# Species-specific secretion of ESX-5 type VII substrates is determined by the linker 2 of EccC_5_

**DOI:** 10.1101/765206

**Authors:** C. M. Bunduc, R. Ummels, W. Bitter, E.N.G. Houben

## Abstract

Type VII secretion systems (T7SSs) are used by mycobacteria to translocate a wide range of effector proteins across their diderm cell envelope. These systems, also known as ESX systems, have crucial roles for the viability and/or virulence of mycobacterial pathogens, including *Mycobacterium tuberculosis* and the fish pathogen *Mycobacterium marinum*. We previously observed species-specificity in the secretion of the PE_PGRS proteins by the ESX-5 system [1], in that the *M. tuberculosis* ESX-5 system was unable to fully complement an *M. marinum esx-5* mutant. In this study, we established that the responsible factor for this is the central membrane ATPase EccC_5_, which has three nucleotide binding domains (NBDs). By creating chimeric *M. marinum*/*M. tuberculosis* EccC_5_ constructs, we observed that PE_PGRS secretion is mediated only in the presence of an EccC_5_ containing the cognate linker 2, irrespective of the origin of the EccC_5_ backbone. This region is responsible for linking the first two NBDs and for keeping the first NBD in an inhibited state. Notably, this region is disordered in a EccC crystal structure and is particularly extended in EccC proteins of the different ESX-5 systems. These results indicate that this region is involved in species-specific substrate recognition and might therefore be an additional substrate recognition site of EccC_5_.

## Introduction

Type VII secretion systems (T7SSs) are crucial virulence determinants for pathogenic mycobacteria such as *M. tuberculosis* [2] and *M. marinum* [3]. Pathogenic mycobacteria can have up to five T7SSs, named ESX-1 to ESX-5 [4], of which three, *i.e.* ESX-1, ESX-3 and ESX-5, have been shown to be functional [1, 5–7]. These secretion systems are paramount for diverse processes, such as the utilization of nutrients and iron or completion of the macrophage infection cycle. In pathogenic mycobacteria, ESX-1 is crucial for intracellular survival by mediating phagosomal membrane rupture [5, 7] and its importance is further substantiated by the fact that the lack of a large part of the *esx-1* gene cluster through the RD1 deletion is the decisive factor in the attenuation of the live vaccine strain *Mycobacterium bovis* BCG [7–9]. Both ESX-3 and ESX-5 systems are essential for *in vitro* growth and have been linked to iron and fatty acid uptake, respectively [1, 6, 10].

In mycobacteria, T7SSs secrete a diverse array of substrates, which includes monomeric as well as heterodimeric protein pairs. The most well-known substrates are the Esx proteins. The mycobacterial Esx proteins are small proteins that form heterodimeric complexes. The first described protein of this family is the ESX-1 substrate EsxA (also named ESAT-6) and its secretion partner EsxB (also called CFP-10). Esx proteins belong to a diverse group of so-called WxG100 proteins, named after their small size of ∼100 residues and a conserved WxG motif located in the turn of the helix-turn-helix structure. For mycobacterial Esx protein pairs, only one of the partner proteins contains the WxG motif, while the other partner protein harbors a C-terminal conserved and critical secretion motif, YxxxD/E [11]. Two major classes of substrates, called the PE and PPE proteins, also form stable heterodimers [12, 13]. The PE protein, named after a conserved proline (P) and glutamic acid (E) motif located N-terminally, forms a conserved ∼110 residues long N-terminal helix-turn-helix structure followed by an YxxxD/E motif, similar to Esx proteins [11]. PPE proteins, named after a similar conserved motif with an extra proline (P) at the N-terminus, have a larger conserved N-terminal domain of ∼180 amino acids, which contains the typical WxG motif located within a turn of the helix-turn-helix structure and additionally consists of a so-called helical-tip domain that is not involved in the dimerization of the PPE with the PE protein. Both PE and PPE proteins can have additional C-termini that are highly variable and might make-up the functional domain of the substrate [14]. Most of the large array of PE and PPE proteins are secreted by the ESX-5 system [1, 15]. ESX-5 is also the most recently evolved mycobacterial T7SS and is present only in the so-called slow-growing mycobacterial cluster, which includes most pathogenic species. A major group of these families that are secreted by the ESX-5 systems are the PE_PGRS proteins, named after the polymorphic <u>G</u>C-rich repetitive <u>s</u>equence motifs in their genes. They have been postulated to be involved in virulence [16–18], whereas other studies implicated their roles in virulence attenuation [19].

The different mycobacterial T7SSs contain a set of conserved components, including two cytosolic and five membrane localized proteins. The cytosolic chaperone EspG has been shown to be involved in substrate recognition [20–23]. EspG interacts specifically with PE/PPE heterodimers and helps to keep these dimers soluble by binding to a hydrophobic patch on the helical-tip domain of the PPE protein [20–22]. By swapping the helical-tip domain of PPE substrates of different systems, the system-specificity of these substrates can be changed [23], showing that this domain is involved in determining system-specificity. Four of the conserved membrane components (EccBCDE) assemble into a hexameric complex of approximately 1.8 MDa, and a first structural image of a T7SS membrane complex, *i.e.* of an ESX-5 membrane complex, was recently provided by negative stain electron microscopy [24, 25]. The dimensions of this complex dictates that it only spans the mycobacterial inner membrane. It therefore remains unknown how substrates are transported across the mycobacterial outer membrane. The fifth conserved and essential membrane component, the subtilisin-like protease mycosin or MycP, interacts only transiently with this complex and is involved in complex stabilization [26]. In addition, mycosins are probably also involved in cleaving specific substrates, although protease activity has only been shown for MycP_1_, that cleaves the ESX-1 substrate EspB upon secretion[27].

A central component of T7SS is EccC, which is a membrane-associated ATPase and most-likely the motor protein of the membrane complex [25]. Importantly, EccC is the only conserved membrane protein in the more distantly related type VII secretion systems of *Firmicutes* [4]. EccC contains two predicted N-terminal transmembrane domains, three nucleotide binding domains (NBDs) and an extra flexible domain of unknown function between de ATPase and transmembrane domains (Fig. 1B, Fig. 1C) [24, 28]. All three NBDs of EccC are part of a family of so-called P-loop NTPases that show strong similarities to FtsK/SpoIIIE proteins, which are proteins that use the energy released from ATP hydrolysis to drive the translocation of macromolecules [29]. Whereas the activity of NBD2 and NBD3 of EccC has been shown to be partially dispensable for secretion, it is NBD1, normally held in an autoinhibited state, that is crucial for T7SS activity [1, 28]. The EccC protein of the ESX-1 system has the unique feature that it is split up in two subunits, *i.e.* EccC_a1_ and EccC_b1_. The C-terminal 7 amino acids of EsxB have been shown by yeast-two-hybrid and pulldown analysis to interact with the EccC_b1_ subunit [11, 28]. Subsequently, a structural analysis of a co-crystal of EccC from *Thermonospora curvata* and a peptide mimicking the C-terminal domain of EsxB revealed that again the last 7 aa of the peptide was bound to NBD3 of EccC [28]. However, substrate-binding to EccC has only been shown for Esx proteins and it therefore remains unclear whether and how the other substrate classes, in particular the major substrate group of PE and PPE proteins, bind to this membrane ATPase. In addition, to what extent EccC is able to recognize substrates in a system-specific fashion is still very poorly understood.

**Figure 1.**
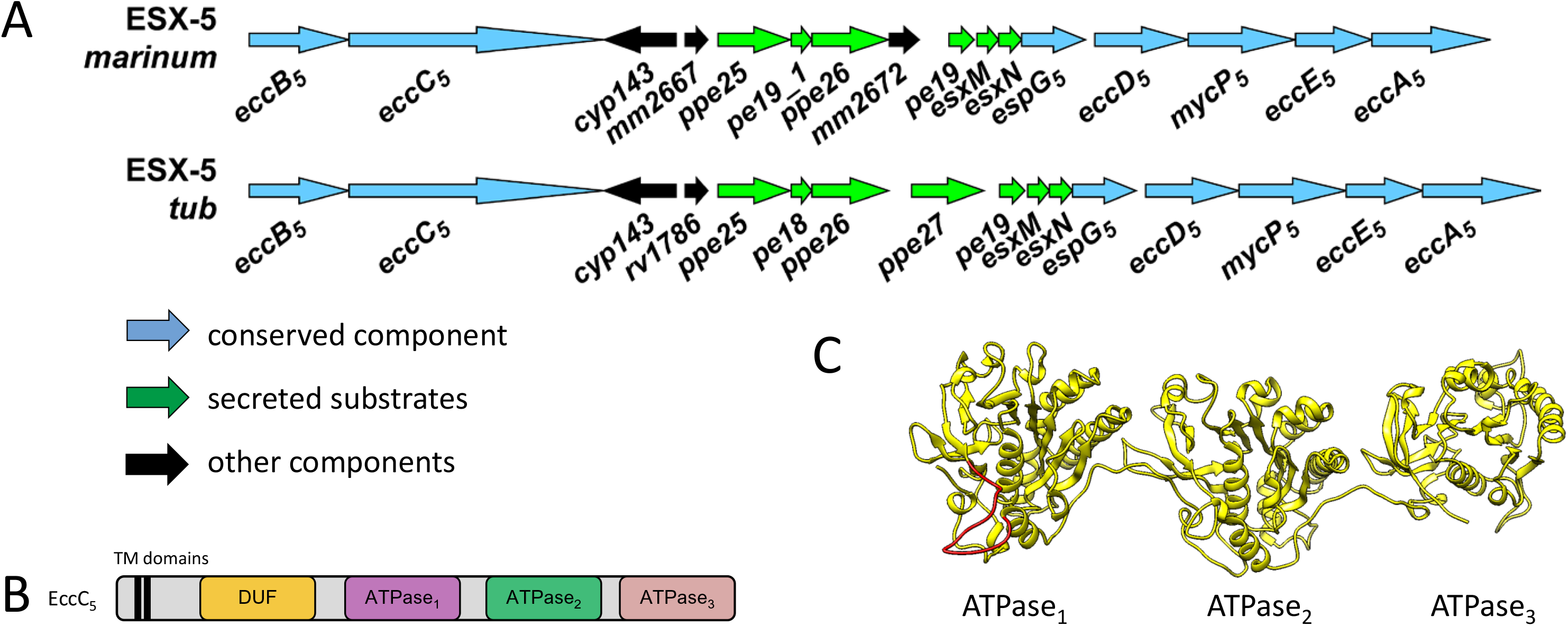
Genetic organization of *esx-5* loci. (A) Genetic organization of the *esx-5* loci in *M. marinum* and in *M. tuberculosis.* (B) General domain architecture of mycobacterial EccC ATPases. The constructs contain: TM – transmembrane domains, DUF – globular flexible domain of unknown function, ATPase – ATP hydrolyzing domain. (C) Structural model of the *M. marinum* EccC_5_ ATPase domains. Generated with Phyre2 (using residues 432 – 1388) with the structure of *T. curvata* EccC as template. In red, linker 2 region missing 65 residues that could not be modeled.

Previous results have shown that the ESX-5 system is essential for growth in the fish pathogen *Mycobacterium marinum* [1]. Strikingly, this essentiality can be circumvented by increasing the permeability of the mycobacterial outer membrane, either by mutating the biosynthesis of the outer membrane lipids phthiocerol dimycocerosates (PDIM) or by introducing an outer membrane porin from *M. smegmatis*, called MspA [1]. As a third option, expression of the homologous *esx-5* operon from *M. tuberculosis* also allowed for the successful deletion of the entire operon in *M. marinum* [1]. Although the ESX-5 system of *M. tuberculosis* is able to mediate growth of the *M. marinum* Δ*esx-5* mutant, the complementation is only partial as the *M. tuberculosis* system is unable to mediate the secretion of many *M. marinum* ESX-5 substrates, most of which are *M. marinum*-specific PE_PGRS proteins [1]. This is unexpected because of the high identity of ∼78% between the two loci (Fig. 1A). As the *M. tuberculosis* system is only partially functional in *M. marinum* we speculated that the observed secretion defects are caused by the fact that many *M. marinum* substrates are not recognized by the *M. tuberculosis* ESX-5 system. To this end, we reasoned that the two conserved components shown to be able to recognize substrates, *i.e.* EspG and/or EccC, could be the cause for this observed system-specific secretion by the ESX-5 system. In this study, we tested this hypothesis by investigating the species-specific roles of EspG and EccC in ESX-5 mediated secretion.

## Results

### EspG_5mtub_ complements secretion of an *M. marinum* Δ*espG_5_*mutant

EspG is a dedicated T7SS chaperone present in four of the five ESX systems and specifically binds PE/PPE proteins in a system-specific fashion. As we hypothesized that the inability of the *M. marinum* Δ*esx-5*::*esx-5_mtub_*to secrete most *M. marinum* PE/PPE substrates is due to the species-specific recognition of these substrates, EspG_5_ was a prime candidate for causing this effect. To test this, we used an Δ*espG_5_*knock-out strain in an *M. marinum* strain that expresses MspA to circumvent the essentiality of ESX-5 for growth [30]. As previously demonstrated, this mutant showed abolished secretion of PE_PGRS proteins, as determined by analyzing their presence in a cell surface fraction extracted by the mild detergent Genapol X-080, as well as EsxN, by analyzing its presence in culture supernatants (Fig. S1). This phenotype could be restored to WT levels of PE_PGRS secretion upon complementation with the *M. marinum espG_5_* gene, although EsxN secretion was only partially restored (Fig. S1). Also the introduction of the *M. tuberculosis espG_5_* gene restored secretion to similar levels (Fig. S1). We therefore conclude that EspG_5mtub_ is fully functional in *M. marinum* and not the cause for the *M. marinum Δesx-5::esx-5_mtub_*species-specific secretion defect of PE_PGRS proteins.

### EccC_5mtub_ complements essentiality but not PE_PGRS secretion in an *M. marinum* Δ*eccC_5_*mutant

In addition to EspG, also the central membrane ATPase EccC_5_ has been shown previously to bind substrates [28, 31]. To examine whether EccC was also responsible for species-specific binding and secretion functions, we checked whether *eccC_5mtub_* is able to rescue the essentiality phenotype of an *M. marinum* Δ*eccC_5_*strain by introducing the same plasmid into a previously described Δ*eccC_5_*::*eccC_5mmar_*strain that lacks MspA [1]. This strain bears an integrative plasmid containing both *eccC_5mmar_* and a hygromycin resistance marker, which was exchanged by the kanamycin resistant plasmid harboring *eccC_5mtub_*. Multiple colonies appeared that showed kanamycin resistance and hygromycin sensitivity, demonstrating the plasmid exchange was successful and therefore that *eccC_5mtub_* is able to mediate growth of an *M. marinum* Δ*eccC_5_* mutant.

Next, we tested the secretion profile of a newly generated Δ*eccC_5_*mutant expressing MspA and also this mutant was complemented with both the *M. marinum* and *M. tuberculosis* versions of *eccC_5_*. As expected, this MspA-expressing *ΔeccC_5_* mutant showed no secretion of PE_PGRS or EsxN substrates (Fig. 2C). While secretion could be restored to WT levels upon complementation with the *M. marinum eccC_5_*gene, introduction of a plasmid containing *eccC_5_* from *M. tuberculosis* showed no visible PE_PGRS secretion in the cell surface enriched protein sample (Fig. 2C). Surprisingly, the culture supernatant fraction of this strain showed only one PE_PGRS protein band, which was similar to the previously observed PE_PGRS secretion phenotype of *M. tuberculosis* (Fig. 2C; Fig. 3)[25]. In addition, no EsxN secretion was observed in this strain. The difference in secretion is surprising, as the overall identity between the two *eccC_5_* orthologs is ∼93%. In addition, the residues present in the pocket of NBD3 that have been predicted to be crucial for Esx interaction [28] are all conserved between the two species. On the other hand, the most C-terminal residues of EsxM, a region that is known to be important for secretion and predicted to be involved in EccC binding, are not well conserved between *M. marinum* and *M. tuberculosis* (Fig S4B). We also checked the colony phenotype for the mutant and the complemented strains. Opposed to the WT and the Δ*eccC_5_*::*eccC_5mmar_*strain, which showed smooth colony morphology and grew as monodispersed cultures, the Δ*eccC_5_* as well as the Δ*eccC_5_*::*eccC_5mtub_*strains grew with a nondispersed phenotype in culture and showed flat and dry colonies on plate (Fig S2). Together, these results show that *eccC_5mtub_* cannot fully complement the *eccC_5_* mutation in *M. marinum* due to species-specific functioning.

**Figure 2.**
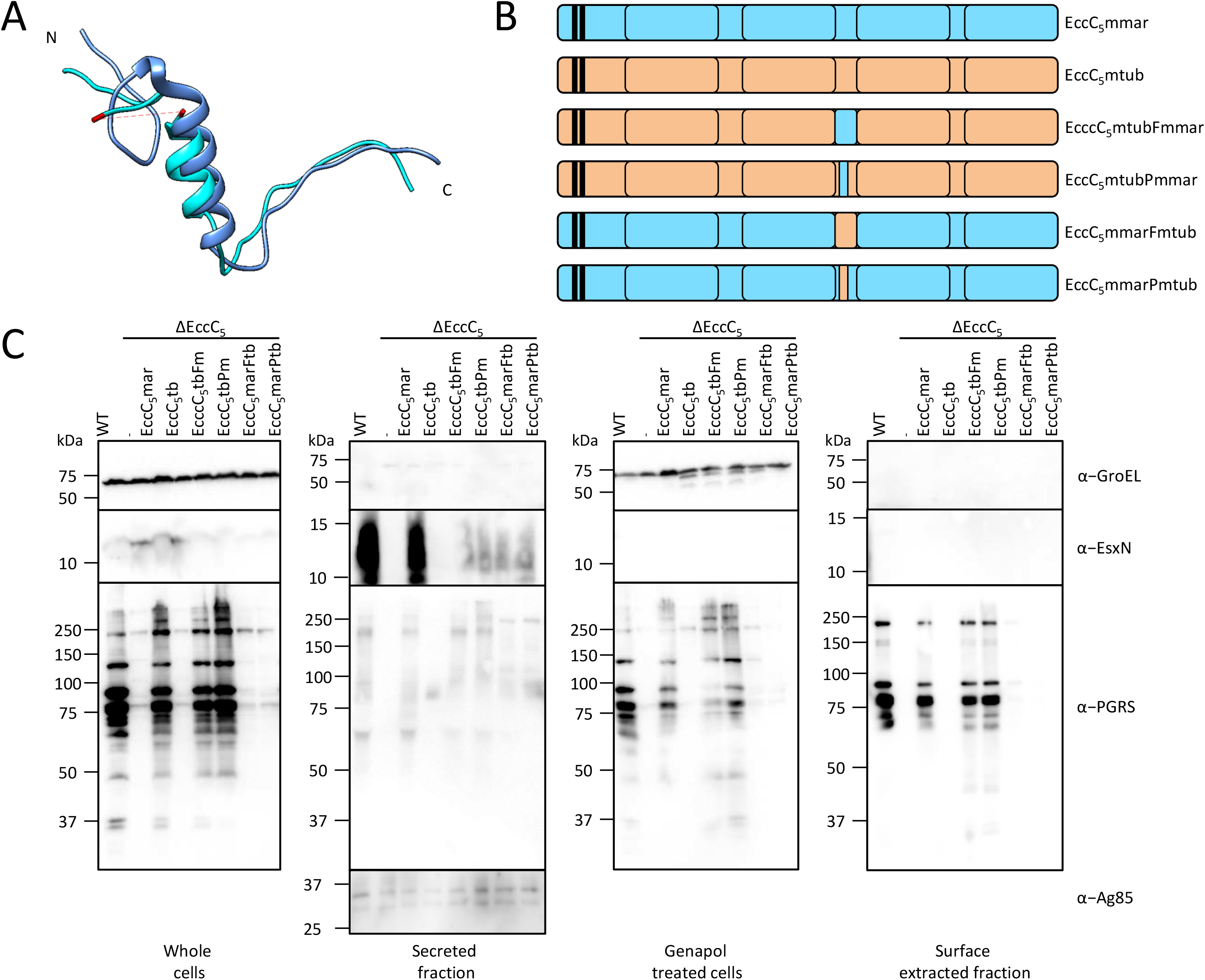
Role of linker 2 in substrate specificity in *M. marinum* ESX-5. (A) *T. curvata* linker 2 in bright blue superimposed with linker 3 in dark blue. In red, the 43 residues that were not resolved in the crystal structure. (B) Schematic overview of *M. marinum* and *M. tuberculosis* EccC_5_ as well as the chimeric constructs used to complement the *eccC_5_* mutants of these species. (C) Immunoblot analysis of the *M. marinum* secretion assay. Δ*eccC_5_* complemented with WT *eccC_5mmar_*, *eccC_5mtub_* and chimeric constructs depicted in B and in Fig S4. Proteins were visualized using antibodies against EsxN and PE_PGRS (ESX-5 substrates), GroEL2 (lysis and whole cell loading control) and Ag85 (secreted fraction loading control).

**Figure 3.**
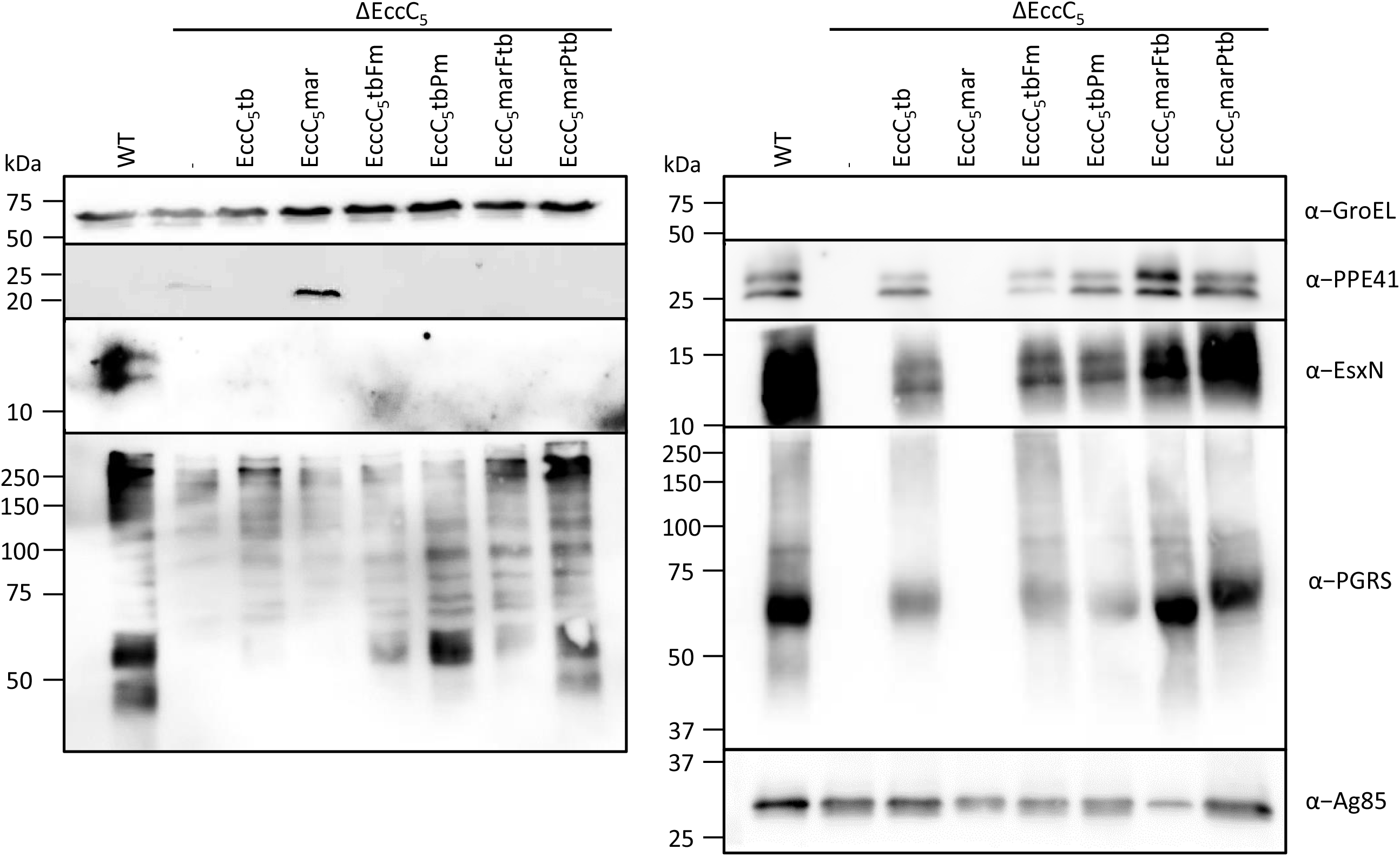
Role of linker 2 in substrate-specificity in *M. tuberculosis* ESX-5. (A) Immunoblot analysis of *M. tuberculosis* secretion assay. EccC_5_ transposon mutant complemented with WT *eccC_5mmar_*, *eccC_5mtub_*and chimeric constructs depicted in Fig. 2B and Fig S4. Proteins were visualized using antibodies against EsxN, PE_PGRS and PPE41 (ESX-5 substrates), GroEL2 (lysis and whole cell loading control) and Ag85 (secreted fraction loading control).

**Figure 4.**
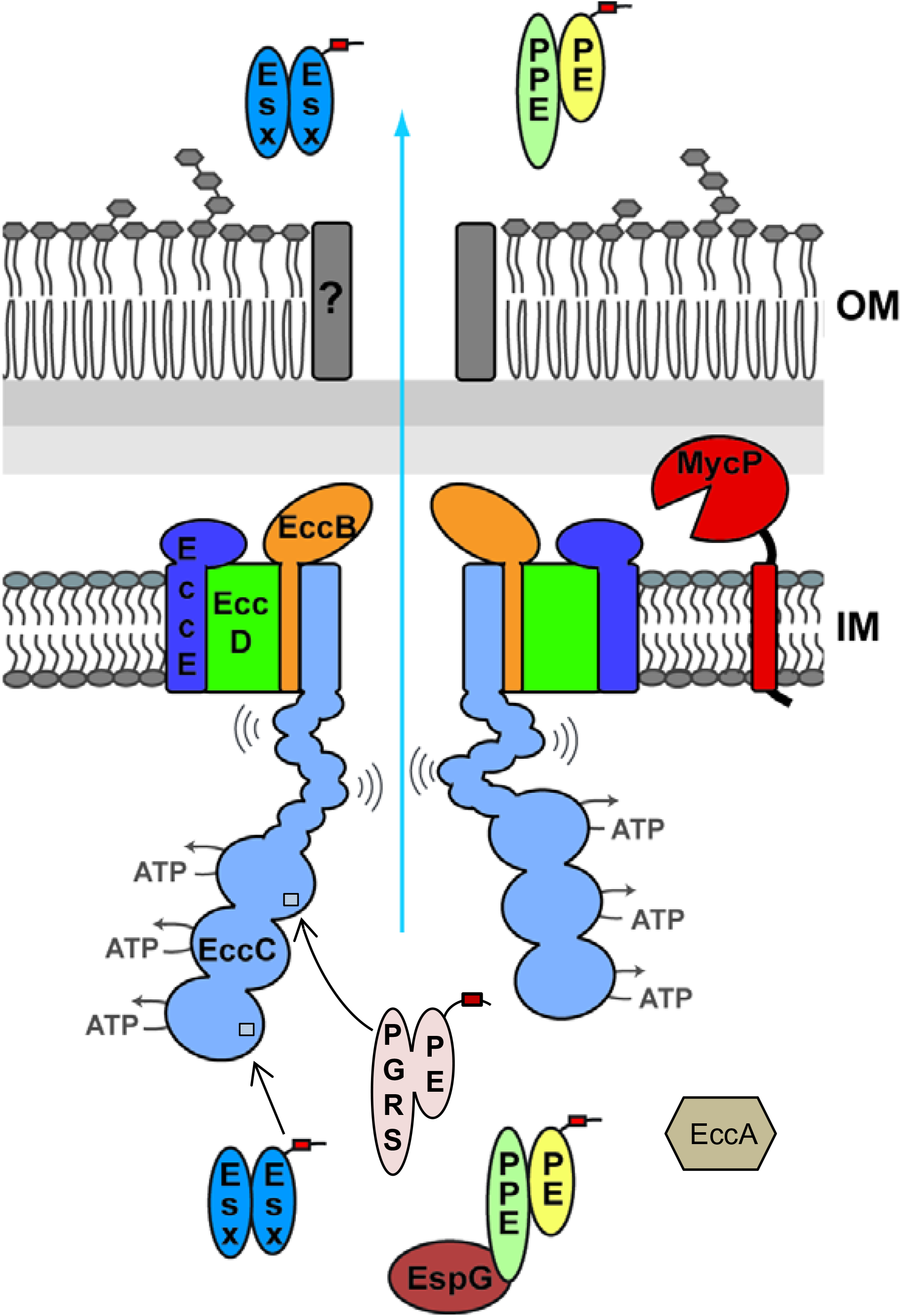
Model for type VII secretion. Substrate recognition occurs at two separate sites on the EccC ATPase. The third NBD interacts with the C-terminus of specific Esx proteins, leading to the multimerization of the soluble domain of EccC. A second site of substrate recognition is located on the first NBD, in close proximity of the linker 2 domain. As the interaction between NBD1 and this linker has an inhibitory effect on the activity of NBD1, substrate binding disrupts this interaction and activates the first ATPase domain. In ESX-5 systems, secretion of PE_PGRS substrates is highly dependent on this domain.

We next investigated whether these secretion defects were due to the unsuccessful incorporation of the EccC_5mtub_ in the ESX-5 membrane complex. We have shown previously that the ESX-5 membrane complex of 1.8 MDa can be observed using BN-PAGE and western blot analysis of DDM solubilized cell envelopes using antibodies against the complex component EccB_5_ [25]. The Δ*eccC_5_* strain showed reduced expression of the ESX-5 membrane components EccB_5_ and EccE_5_ and membrane complex formation was abrogated (Fig. S3A). Complex formation was restored upon complementation with either the *M. marinum* or *M. tuberculosis eccC_5_* containing plasmid (Fig. S3A). Similarly, expression of the EccB_5_ and EccE_5_ components were restored to WT levels (Fig S3B). We therefore conclude that the lack of PE_PGRS secretion by the *M. marinumΔeccC_5_::eccC_5mtub_* was not caused by any defect in the assembly of the ESX-5 membrane complex. Because the secretion phenotype of the Δ*eccC_5_*::*eccC_5mtub_*strain was similar to that of the whole *esx-5_mtub_* complementation strain, we conclude that EccC is the key component responsible for this distinct secretion defect.

### EccC_5_ linker 2 domain is involved in species-specific secretion in *M. marinum*

Although EccC_5mtub_ is properly integrated in the ESX-5 membrane complex and rescues essentiality of an *M. marinum ΔeccC_5_*mutant, this did not restore secretion. A sequence alignment of the EccC_5_ genes of *M. marinum* and *M. tuberculosis* showed high overall conservation, but also revealed some variations (Fig. S4A, Table S1). Aligning the two genes with the sequence of the *T. curvata* EccC, which crystal structure has been previously solved [28], revealed that the amino acids and motifs that were shown to be important for ATPase activity and substrate-binding are highly conserved (Fig. S4, Table S1). In particular, the arginine present in pocket1 of NBD1 (R563_mtub_ and R564_mmar_) as well as its two interacting residues in the linker 2 region, *i.e.* tryptophan (W810_mtub_ and W807_mmar_) and glutamine (Q811_mtub_ and Q808_mmar_ (L763 in *T. curvata*)), which form the major interaction between this linker region and NBD1. Similarly, the interacting amino acids lining the substrate binding pocket on NBD3, *i.e.* E1237, L1253, I1282 for *eccC_5mtub_*and E1234, L1250, I1279 for *eccC_5mmar_* (I1163, I1179 and L1208 in the *T. curvata* system), are all conserved (Table S1). However, alignment analysis also revealed key differences, particularly in the linker 2 region between NBD1 and NBD2 (Fig. S4). For EccC of *T. curvata* this linker 2 domain was shown to be crucial in keeping the critical first NBD in an inhibited state [28]. Based on this, it has been speculated that a yet unknown event might lead to the displacement of linker 2 from the pocket of NBD1, allosterically regulating its activity [28]. Interestingly, a significant part (41 residues) of this linker 2 domain was disordered and therefore not present in the crystal structure of EccC of *T. curvata* (Fig. 2A). This disordered region also revealed the lowest sequence identity between the *eccC* genes of *M. tuberculosis* and *M. marinum*. As compared to the *T. curvata* EccC protein, the linker 2 region is considerably larger for EccC_5_ proteins, with an additional 31 residues for EccC_5mtub_ and 27 residues for EccC_5mmar_.

Taking these observations into consideration, we reasoned that this disordered region in linker 2 might play a crucial role in the (in)activation of NBD1 through regulating substrate binding and/or specificity. In order to test this, we made 2 chimeric *eccC_5_* constructs where the backbone originates from *M. tuberculosis* and the linker 2 region from *M. marinum*. The linker 2 portion covered either the entire region after NBD1 until just after the two amino acids WQ that interact with the pocket1 (named full – EccC_5mtub_F_mmar_) or only a small portion of the linker 2 that shows the most sequence divergence between the two, and that also aligns broadly with the disordered region (named partial – EccC_5mtub_P_mmar_)(Fig. 2B, Fig S4A). Both constructs could rescue the essentiality of the *M. marinum eccC_5_* knockout in the absence of MspA. Subsequently, we examined the expression of ESX-5 membrane components and the presence of the ESX-5 membrane complex by BN-PAGE immunoblot analysis, which showed that both proteins were incorporated (Fig 3 SAB). Finally, while expression of the original EccC_5mtub_ in the *eccC_5_*mutant resulted in flat and dry colonies, this phenotype was reversed to the WT situation upon exchange of the linker 2 in the EccC_5mtub_F_mmar_ or EccC_5mtub_P_mmar_ plasmids (Fig. S2).

With these constructs we next checked if the secretion defect caused by the EccC_5mtub_ complementation could be restored. Remarkably, although EsxN was not fully restored, secretion of PE_PGRS proteins was restored to WT levels with both full and partial swapped constructs (Fig. 2C). This is intriguing, as the swapped region in the partial construct has only 19 amino acid difference as compared to the WT *eccC_5mtub_* gene. These data confirmed our hypothesis that linker 2 is involved in substrate specificity and/or (in)activation of NBD1.

Next, we wondered if conversely, we could repress secretion of the Δ*eccC_5mmar_*::*eccC_5mmar_* complementation by swapping the linker 2 of EccC_5mmar_ with that of EccC_5mtub_. Importantly, although the linker 2 region originated from EccC_5mtub_ the rest of the gene was *M. marinum* WT, thus keeping all other (un)known potential interaction sites. These constructs were named EccC_5mmar_F_mtub_ and EccC_5mmar_P_mtub_ (Fig.2B, Fig. S4A). Although only 25 residues for the full linker 2, or 19 residues for partial linker 2 were different, PE_PGRS protein secretion for the new chimeric constructs was completely abolished, thus substantiating our initial finding. Conversely, EsxN was present in the supernatant, but only in low amounts, similar to the reciprocal chimeric constructs. Both constructs could rescue the essentiality of the *M. marinum* Δ*eccC_5_* mutant and showed a somewhat intermediate phenotype between the smooth WT colonies and the rough and dry colonies of the Δ*eccC_5_* and Δ*eccC_5_::eccC_5mtub_*. From this, we conclude that the linker 2 domain of EccC_5_ is involved in species-specific secretion of PE_PGRS substrates. In addition, as EsxN secretion is only partially recovered by all chimeric constructs, the optimal secretion of EsxN is not only dependent on NBD3, but is regulated by multiple domains or interactions with EccC.

### EccC_5_ linker 2 is involved in substrate specificity in *M. tuberculosis*

With the drastic effects upon exchanging only a small part of the linker 2 of EccC_5_ observed in *M. marinum*, we questioned whether this phenotype is more general. To test if our results could also apply to other species, we used a previously described *M. tuberculosis eccC_5_*transposon mutant and introduced the same chimeric constructs as analyzed in *M. marinum* (Fig. 2B, Fig S4A)[25].

The different constructs were efficiently expressed in the *M. tuberculosis* mutant strain (Fig. S5). In addition, while expression of EccB_5_ was strongly affected in the mutant strain, its expression was restored to WT levels in the presence of all constructs. From this, we conclude that the chimeric EccC proteins were again able to stabilize other components of the ESX-5 membrane complex, suggesting these constructs were properly integrated in the membrane complex (Fig. S5). In this case, membrane complex formation was not checked via BN-PAGE as biosafety measurements prohibited the membrane isolation procedures required for this.

Similar to *M. marinum*, in *M. tuberculosis* secretion of PE_PGRS proteins and EsxN was completely abolished in the absence of *eccC_5_*, and this could be restored by the WT *M. tuberculosis* gene, albeit to a slightly lower level (Fig. 3). Intriguingly, the *M. marinum* gene was not able to complement both PE_PGRS and EsxN secretion. Upon exchanging the linker 2 of the *M. marinum* protein with that originating from *M. tuberculosis* EccC_5_, secretion was restored to WT levels. However, the chimeric EccC_5mtub_ with the linker 2 from EccC_5mmar_ also showed PE_PGRS and EsxN secretion restored back to WT levels (Fig. 3). From this we can conclude that linker 2 domain has a similar role in *M. tuberculosis*, as described for *M. marinum* ESX-5.

## Discussion

ESX-5 is the most recently evolved T7SS in mycobacteria and is responsible for the secretion of most PE/PPE proteins, amongst them most or even all members of the large family of PE_PGRS proteins. The ESX-5 system is essential for cell viability of *M. marinum*, which is linked to outer membrane permeability [1]. Previous studies have shown that introduction of the ESX-5 system of *M. tuberculosis* is able to take over the essential role of the ESX-5 system in *M. marinum*. However, this system is only marginally able to restore secretion of *M. marinum* ESX-5 dependent substrates, suggesting that substrate recognition is at least partially species-specific. This study revealed that a highly specific domain in the central membrane ATPase of the ESX-5 system, *i.e.* linker 2 of EccC_5_, is the determining factor for the specifies-specific secretion of PE_PGRS proteins.

To identify the specific component in the observed species-specific secretion of PE_PGRS proteins, we used individual *esx-5* component mutants and complemented these with the corresponding gene from either *M. marinum* or *M. tuberculosis*. Because the *M. marinum* Δ*esx::esx-5_mtub_*showed no secretion of PE_PGRS proteins, we decided to use this as a readout for functionality. Additionally, by assessing PE_PGRS protein secretion, we could monitor a whole set of substrates and not just individual proteins.

As EspG has been shown to bind PE/PPE heterodimers and the corresponding EspG binding domain is a determinant factor in the system-specific secretion of these substrates, this was our most prominent candidate. However, Δ*espG_5_* complementations did not show any marked difference in secretion between EspG_5_ of *M. marinum* and *M. tuberculosis*. As the conservation between the two genes is very high, ∼97% identity, probably both of them can serve each other’s function in the opposite species.

The only other component known to recognize substrates is EccC, although only Esx substrates have been shown to interact with this central ATPase component[28, 31]. Complementation with *eccC_5mtub_*of *M. marinum* Δ*eccC_5_* abolished the presence of PE_PGRS proteins on the cell surface, while in the supernatant only a single PE_PGRS band remained present. Importantly, localization of PE_PGRS proteins to the cell surface and to the culture supernatants could be fully restored by introducing only a small part of the linker 2 region, between NBD1 and NBD2, of the *M. marinum* EccC into the *M. tuberculosis* protein. EsxN secretion is recovered only partially under this condition. Similarly, introducing the linker 2 region of EccC_5mtub_ into the *M. marinum* protein was not able to restore cell-surface localization of PE_PGRS proteins, while EsxN secretion remains at a similar basal level. The absence is also linked to a distinct flat and dry colony morphology. Interestingly, this EccC_5_ mmar-mtub chimera was able to mediate secretion of PE_PGRS proteins into the culture supernatant, although the band pattern is distinct from that of the WT strain. Significantly, our results were similarly replicable in *M. tuberculosis*. Notably, surface-associated PE_PGRS proteins are not extractable in this species [19, 25], and in our *M. marinum* experiments, this subset of substrates is completely dependent on the cognate linker 2. As such, due to technical limitations, we are not able to analyze the impact of our EccC_5_ chimeras on the surface localization of PE_PGRS proteins in this species. However, analysis of the PE_PGRS substrates in the secreted fraction shows an identical phenotype. Whereas secretion analysis of an *eccC_5_* mutant shows abolished secretion of PE_PGRS and EsxN in either species, complementation with the native gene restores this back to WT levels. Similarly, introduction of a plasmid containing *eccC_5_* from the opposite species abolishes secretion to the culture supernatant completely. However, expression of an EccC_5_ chimera which contains the native linker 2 and the backbone of the opposite species, restores secretion in both *M. marinum* and *M. tuberculosis*. This further strengthens the hypothesis that the EccC_5_ linker 2 domain is involved in determining substrate-specificity for the secretion of PE_PGRS substrates.

Although the interface between NBD1 and NBD2 is similar to that between NBD2 and NBD3, the interdomain linkers are variable. Both interdomain linkers form a main α-helix which mimic the C-terminal tail of EsxB like proteins that binds on NBD3. However, immediately N-terminal from of this α-helix, there is a region of variability between linker 2 and linker 3 in sequence and size, *i.e.* this region is significantly larger in the linker 2 interdomain. Highly intriguing is that a large part of this variable region of linker 2 is disordered in the only available structure containing all three NBDs of an EccC homologue from *T. curvata*, suggesting flexibility (Fig 2A, S.Fig. 6AB, Table 1). Sequence alignments show that this is also the most variable region between the two *eccC_5_*genes of *M. tuberculosis* and *M. marinum* (Table S1). Aligning the EccC ATPases from all five mycobacterial ESX systems shows this region of the linker 2 domain to be extended in ESX-2 and ESX-5 systems, the most recent systems (Table S2). ESX-5 most-likely evolved through a duplication of the *esx-2* cluster [32], and this duplication event is followed by the vast expansion of *pe* and *ppe* genes, most notable the most-recently evolved PE_PGRS proteins and the so-called PPE_MPTR proteins, of which at least a major portion are secretion by ESX-5 [15, 16]. Concurrently, different ESX-5 dependent PE/PPE heterodimers from *M. tuberculosis* or *M. marinum* showed no notable secretion difference in *M. marinum* Δ*eccC_5_::eccC_5mmar_* or Δ*eccC_5_::eccC_5mtub_*(data not shown). As previous mass spectrometry analysis of an Δ*esx-5::esx-5_mtub_*strain showed that most PE/PPE proteins are not secreted by the *M. tuberculosis* system [1], it is tempting to speculate that the extension of the linker 2 domain has co-evolved with the expansion of PE/PPE proteins to allow the recognition of this vast group of substrates.

Importantly, the secretion of EsxN is also species-specific, as both the *M. marinum* and *M. tuberculosis eccC_5_* mutants could only be restored to WT levels upon complementation with the cognate *eccC_5_*gene. This is in contrast to the conservation found in the putative binding pocket of Esx substrates on NBD3. Notably, EsxM, *i.e.* the partner protein of EsxN carrying the predicted C-terminal secretion signal, has a stop codon in *M. tuberculosis* and both EsxN and EsxM have multiple highly homologous paralogs in *M. marinum* and *M. tuberculosis*, which probably have redundant functions. Although there is high conservation between the different EsxM homologs from *M. marinum* and *M. tuberculosis*, the most C-terminal 4 amino acids are divergent. As in the *T. curvata* system the last 7 amino acids of EsxB are involved in EccC-binding [28], this provides an explanation for the species-specific secretion of EsxN. However, our observation that all chimeric EccC_5_ proteins with exchanged linker 2 regions, irrespective of the origin of NBD3, showed intermediate levels of EsxN secretion in *M. marinum*, and full secretion in *M. tuberculosis* is more difficult to explain. As T7SS substrates have been shown to be inter-dependent on each other for secretion [19, 33, 34], it might be possible that the required PE/PPE protein(s) and EsxN are secreted in a concerted fashion. While substrate-interdependency for secretion is a not yet understood phenomenon in T7SS, these data suggest that secretion-specificity of Esx heterodimers is not only dependent on the interaction with its cognate EccC NBD3.

Based on our current results and previous data we propose a working model for substrate-recognition by EccC in mycobacterial T7SSs. In this model the EccBCDE membrane complex is a stable complex in the mycobacterial inner membrane. Without substrate-binding, EccC is hexameric via its transmembrane regions, while its cytosolic domain is highly flexible through the extra N-terminal domain [24]. EccC is kept inactive through the pocket1/linker 2 connection while interaction of the C-terminal NBD3 with the C-terminal signal sequence of specific (Esx) substrates drives multimerization [28]. Although still inactive, multimerization of EccC should be aided also by a conserved arginine residue (“R finger”) which completes the active site of the neighboring ATPase [28]. The final secretion activation step takes place upon linker 2 displacement from pocket 1, which at least in the mycobacterial ESX-5 systems, is triggered by binding of PE/PPE substrates or a yet unknown chaperone for these proteins. While the role of linker 2 of EccC in keeping the highly important NBD1 in an inactive state has been previously described, we present here a new role in substrate-specificity for this domain of EccC ATPases.

## Materials and methods

### Bacterial strains and culture conditions

*M. marinum* M and *M. tuberculosis* CDC155 were used for all experiments involving EccC_5_ and *M. marinum* E11 was used for the EspG_5_ analysis. *M. marinum* was grown at 30°C on 7H10 agar with 10% Middlebrook OADC (BD Biosciences) or in 7H9 liquid medium with 10% Middlebrook ADC and 0.05% Tween 80 (Merck). *M. tuberculosis* was grown at 37°C under similar conditions. Culture media was supplemented with the necessary antibiotics at the following concentrations: kanamycin, 25 µg/ml; hygromycin, 50µg/ml; streptomycin, 30 µg/ml.

### Molecular Cloning

All cloning was performed in *Escherichia coli* DH5α, with restriction enzymes from New English Biolabs and PCR amplifications with Iproof, BioRad. Difficult ligations or ligations that included more than two fragments were performed with In-Fusion, TakaraBio, with 15 bp homologies coded in the primer sequence. The *esxM-esxN-espG_5_* region or *eccC_5_*was amplified from *M. tuberculosis* H37Rv genomic DNA using anchored primers 1 and 2 or 3 and 4, respectively (XmnI, HindIII, Supplementary table 4) and ligated in two modified pMV361 vectors coding for Hyg^r^ or Kan^r^ [1] using XmnI and HindIII, resulting in the plasmids pMV-EspG_5mtub_-hyg^r^ and pMV-EccC_5mtub_kan^r^. Primers 5 and 6 were used to PCR amplify the partial linker 2 domain (between F744 and V786) from the pMV-EccC_5mmar_ plasmid. This PCR product and the pMV-EccC_5mtub_ plasmid were both cut with FspAI and MunI and ligated, resulting in the plasmid pMV-EccC_5mtub_P_mmar_.

Due to a lack of proper restriction sites plasmid pMV-EccC_5mtub_ was cut with SfiI and HindIII, removing the linker 2 domain but also NBD2 and NBD3. Primers 7 and 8 were used to amplify the full linker 2 domain of EccC_5mmar_ (between residues A680 and D819) and primers 9 and 10 were used to PCR amplify the C-terminal part of the EccC_5mtub_ gene, downstream of the linker 2, including NBD2 and NBD3. Primers encoded for 15bp homologies and both PCR products and the cut backbone were ligated via In-Fusion cloning, resulting in plasmid pMVEccC_5mtub_F_mmar_.

pMVEccC_5mmar_F_mtub_ and pMVEccC_5mmar_P_mtub_ were cloned in a similar fashion. Due to a lack of proper restriction sites, these plasmids were cloned from scratch using In-Fusion cloning. The EccC_5mmar_ region upstream of the linker 2 domain was amplified from the pMV-EccC_5mmar_ plasmid with primers 11 and 12 for the full domain (PCR1) and primers 11 and 17 for the partial domain (PCR2). The EccC_5mtub_ linker 2 domain was amplified from the corresponding plasmid with primers 13 and 14 for the full (between residues A679 and D822 – PCR3) and 18 and 19 for the partial domain (between residues F743 and T789 – PCR4). The EccC_5mmar_ region downstream of the linker 2 domain was PCR amplified from pMV-EccC_5mmar_ with primers 15 and 16 for the full domain – PCR5 – and primers 16 and 20 for the partial domain – PCR6. A pMV-kan^r^ plasmid cut with the XmnI and HindIII restriction sites was In-Fusion ligated with PCR products 1+3+5 to result in the plasmid pMV-EccC_5mmar_F_mtub_ and with PCR products 2+4+6 to result in the plasmid pMV-EccC_5mmar_P_mtub_.

### Protein secretion and western blot analysis

For protein secretion, *M. marinum* strains were grown in 7H9 liquid medium with 10% Middlebrook ADC, 0.05% Tween 80 and appropriate antibiotics until mid-log phase. Cells were harvested, washed and inoculated at an 0.4-0.5 OD_600_ in 7H9 liquid medium with 0.2% dextrose, 0.2% glycerol, 0.05% Tween 80 and appropriate antibiotics. After overnight growth, cells were pelleted at an OD_600_ of 0.8-Supernatants were passed through an 0.2 µm filter and precipitated with trichloroacetic acid (TCA)(culture supernatant fraction). Cell pellets were split in two and half was treated with 0.5% Genapol x80 (Fluka) for 30 minutes, head over head at room temperature, after which cells were spun down and supernatant was collected (Genapol surface extracted fraction and Genapol trated cells). Both whole cell samples, treated or not with Genapol-x80 were lysed by bead beating. SDS loading buffer was added and samples were boiled and loaded on SDS-PAGE gels (10%-16%, depending on the size of the proteins of interest), transferred to nitrocellulose membranes and stained with appropriate antibodies. For experiments involving *M. tuberculosis*, the procedure was similar but cells and culture supernatants were heat inactivated for 30 minutes at 80°C after harvesting. The antibodies that were used are anti-GroEL (Cs44; John Belisle, NIH, Bethesda, MD, USA), anti-EsxN *M. marinum* (Mtb9.9a), anti-PE_PGRS [35], anti-Ag85, anti-PPE41 [35], anti-EspG_5_ [25], anti-EccB_5_ [25], anti-EccC_5_ [25], anti-EccE_5_ [25], anti-FtsH [25],

### Cell envelope isolation

For cell envelope isolations, *M. marinum* was grown in liquid 7H9 media with 10% Middlebrook ADC, 0.05% Tween 80 and appropriate antibiotics to an OD_600_ of 1.2-1.5. Cells were washed in PBS and resuspended in CE buffer (20 mM Tris-HCl, 300 mM NaCl and 10% glycerol). Cells were lysed by passed through a One-Shot Cell disruptor (Constant Systems Ltd.) and unbroken cells were pelleted at 5000xg. Cell envelopes (CE) were separated from the soluble fraction by ultracentrifugation at 150.000xg for 90 minutes. After ultracentrifugation, supernatant was discarded, pellets were washed in CE buffer and resuspended in CE buffer.

### BN-PAGE

For BN-PAGE analysis of membrane complexes, cell envelopes were solubilized with 0.25% DDM for one hour at 4°C. Nonsolubilized material was pelleted by centrifugation at 100.000xg for 20 minutes at 4°C. NativePage 5% G-250 Sample Additive (Invitrogen) was added to the resulting supernatant fraction and samples were run on a 3-12% NativePage Bis-Tris Protein Gel (Invitrogen). Gels were blotted to a PVDF membrane and stained with appropriate antibodies.

## Figures

**Supplementary figure 1.**
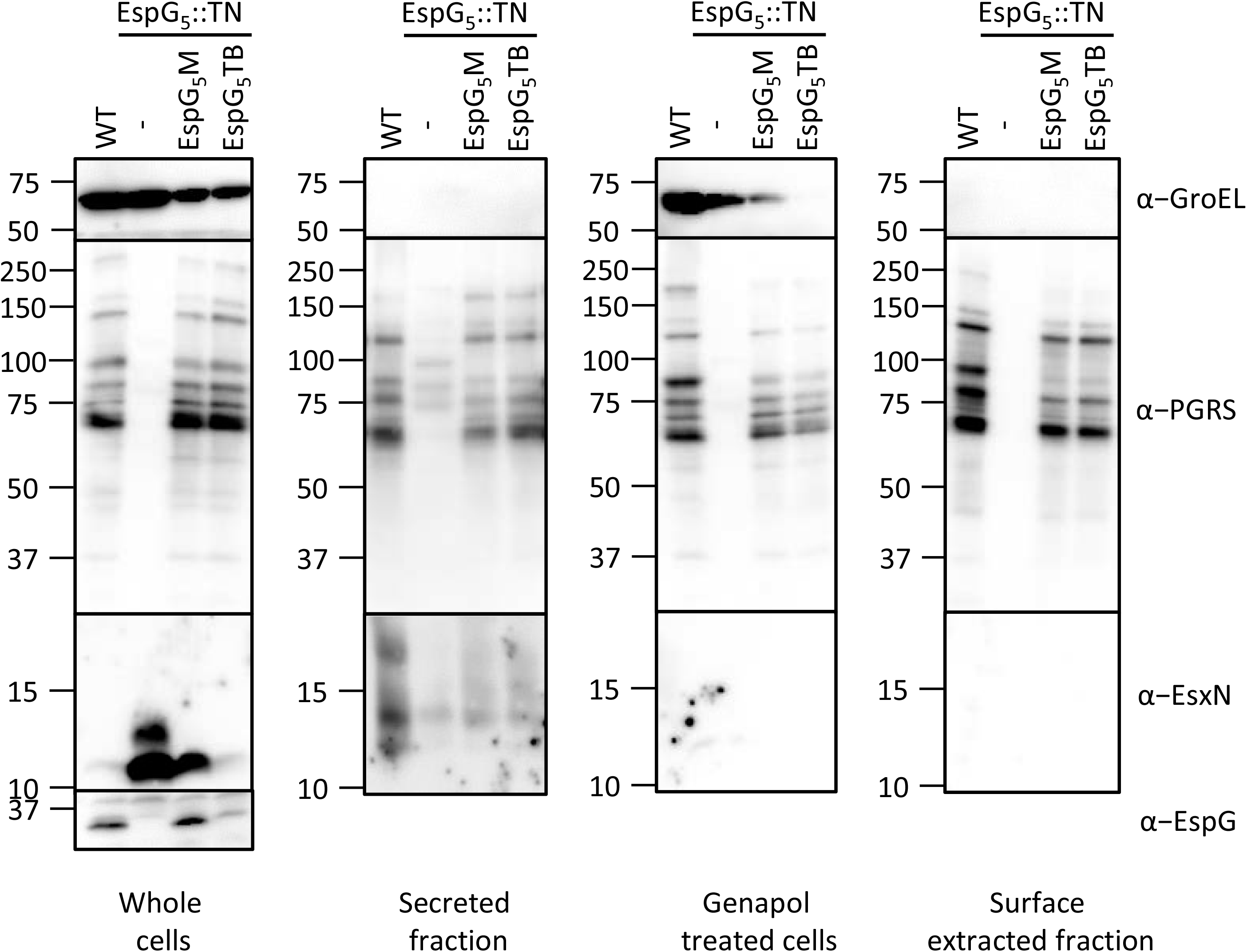
EspG_5mtub_ complements secretion of a *M. marinum* Δ*espG_5_*. Secretion analysis of *M. marinum* WT, *ΔespG_5_* and *ΔespG_5_*complemented with WT *espG_5mmar_* or *espG_5mtub_*. Proteins were visualized using antibodies against EsxN and PE_PGRS (ESX-5 substrates), GroEL2 (lysis and whole cell loading control) and EspG_5_.

**Supplementary figure 2.**
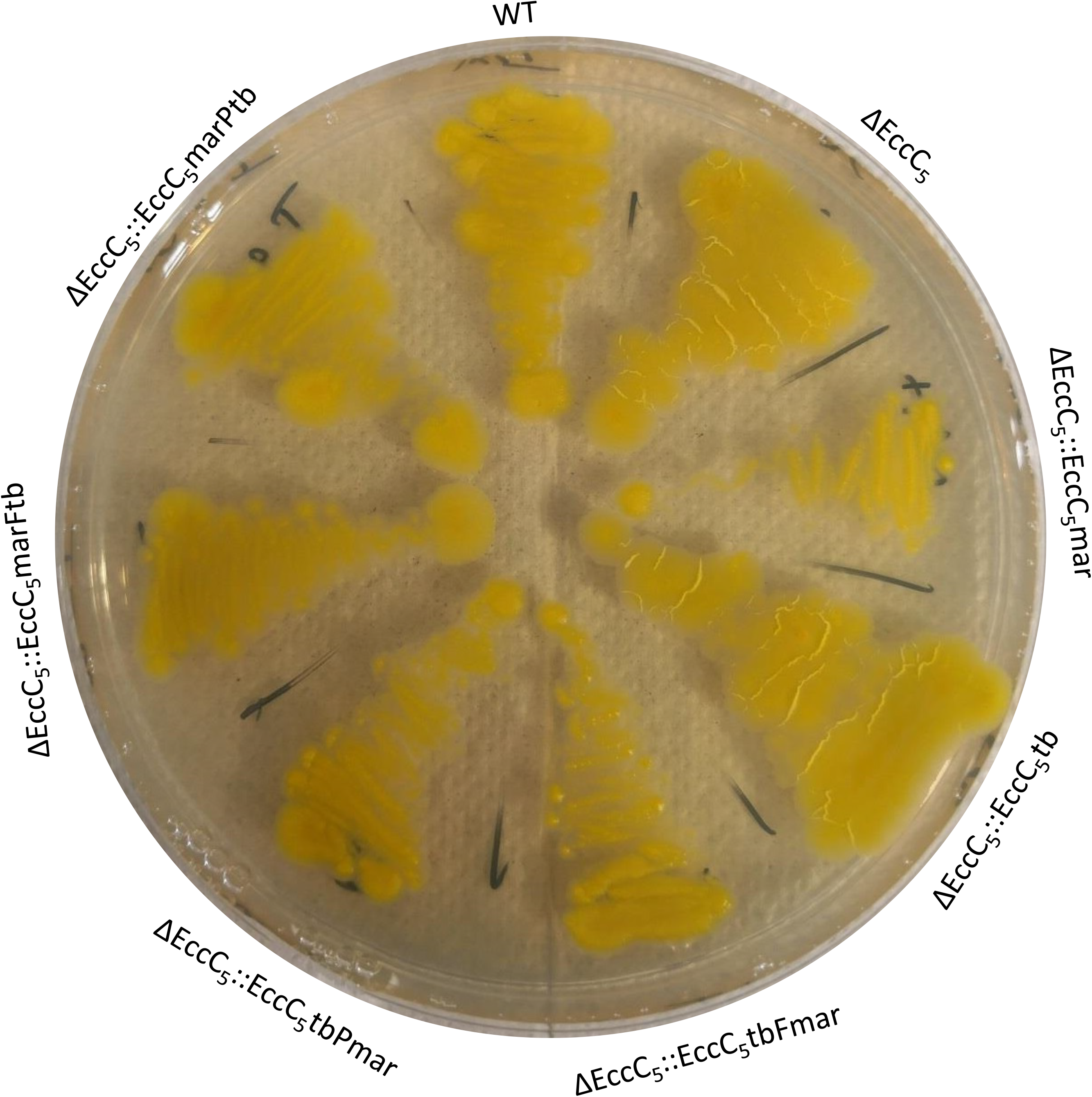
Linker 2 of EccC_5mtub_ affects colony morphology in *M. marinum*. Colony morphology of *M. marinum* WT, *ΔeccC_5_* and *ΔeccC_5_*complemented with constructs depicted in Fig 2B and Fig S4. Strains were streaked on 7H10 agar media supplemented with 10% Middlebrook OADC (BD Biosciences).

**Supplementary figure 3.**
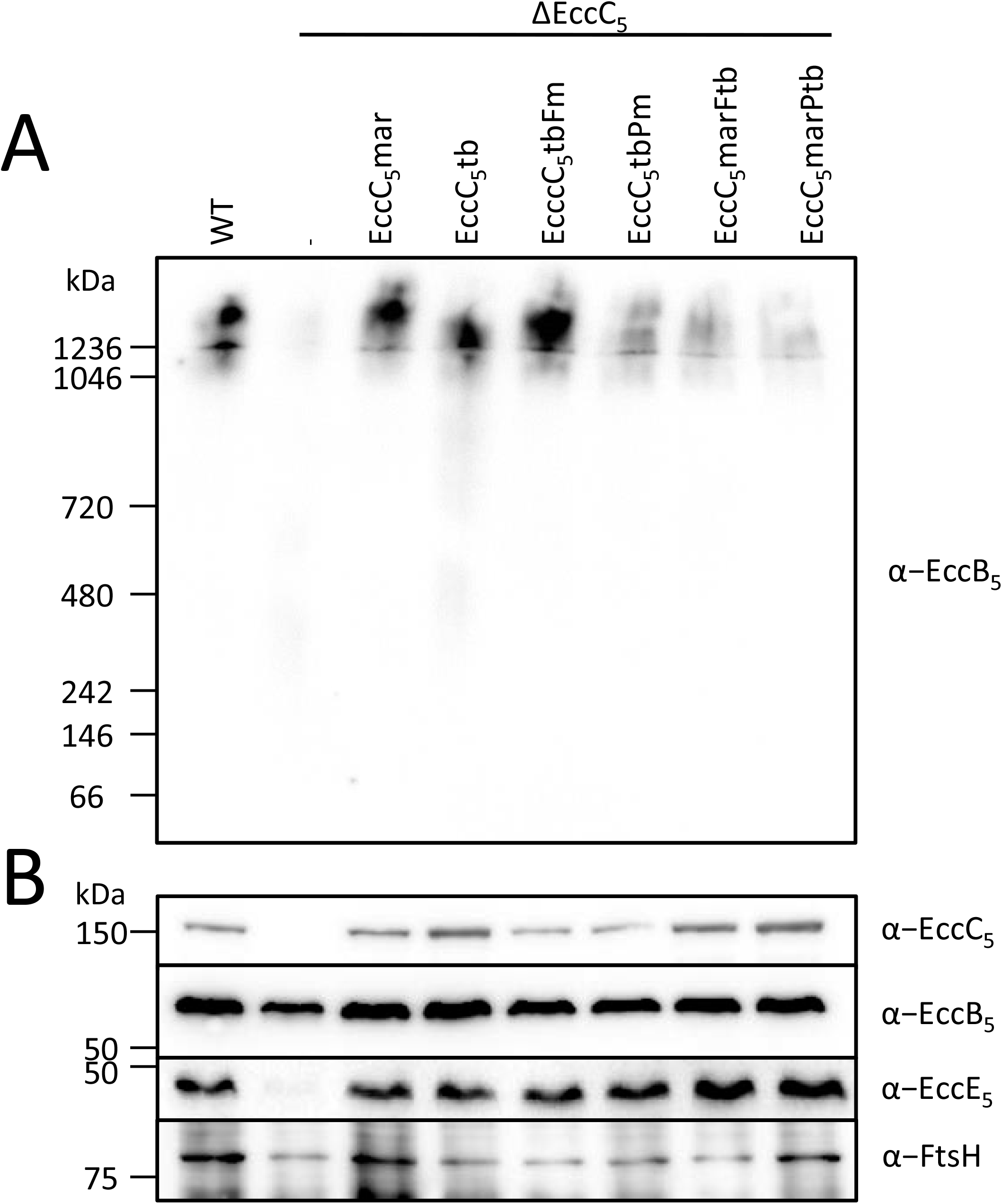
ESX-5 membrane complex analysis. (A) BN-PAGE immunoblot analysis of DDM solubilized cell envelope fractions of *M. marinum* WT, *ΔeccC_5_* and *ΔeccC_5_* complemented with constructs depicted in Fig 2B and Fig S4. (B) Analysis of the same set of samples as in A, but under denaturing conditions. Proteins were visualized using antibodies against EccB_5_, EccC_5_, EccE_5_ (membrane proteins forming the ESX-5 membrane complex) and FtsH (membrane protein loading control).

**Supplementary figure 4.**
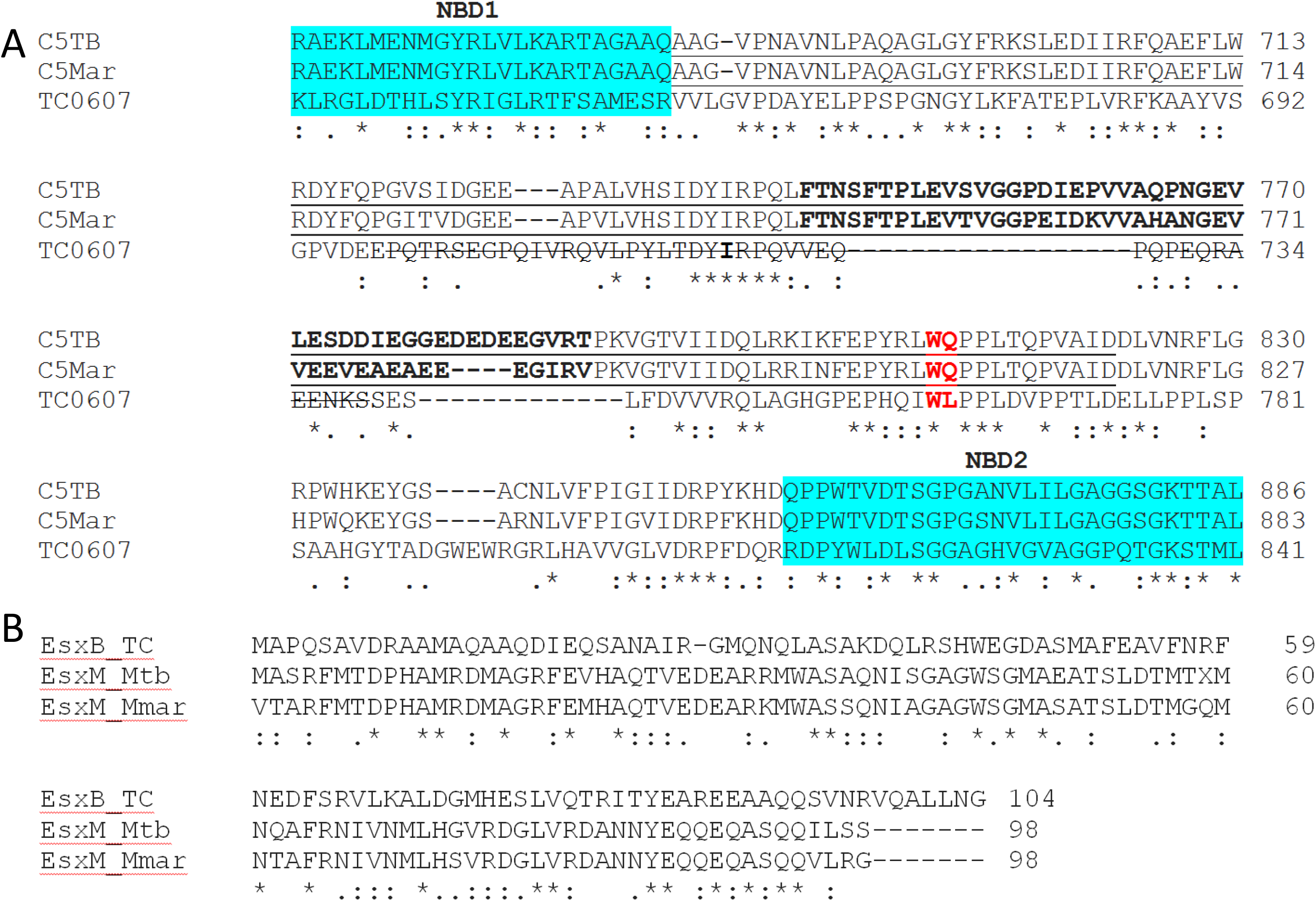
Linker 2 sequence divergence. (A) Alignment of linker 2 region from *M. tuberculosis* EccC_5_, *M. marinum* EccC_5_ and *T. curvata* EccC. In blue the C-terminus of NBD1 and N-terminus of NBD2. In bold, the sequence swapped between constructs EccC_5mmar_P_mtub_ and EccC_5mtub_P_mmar_ (partial linker 2 swap). Underlined, the sequence swapped between constructs EccC_5mmar_F_mtub_ and EccC_5mtub_F_mmar_ (full linker 2 swap). The missing structural feature of the linker 2 in the *T. curvata* EccC crystal structure is striked through. (B) Sequence alignment between EsxB of *T. curvata* and EsxM from *M. marinum* and *M. tuberculosis*.

**Supplementary figure 5.**
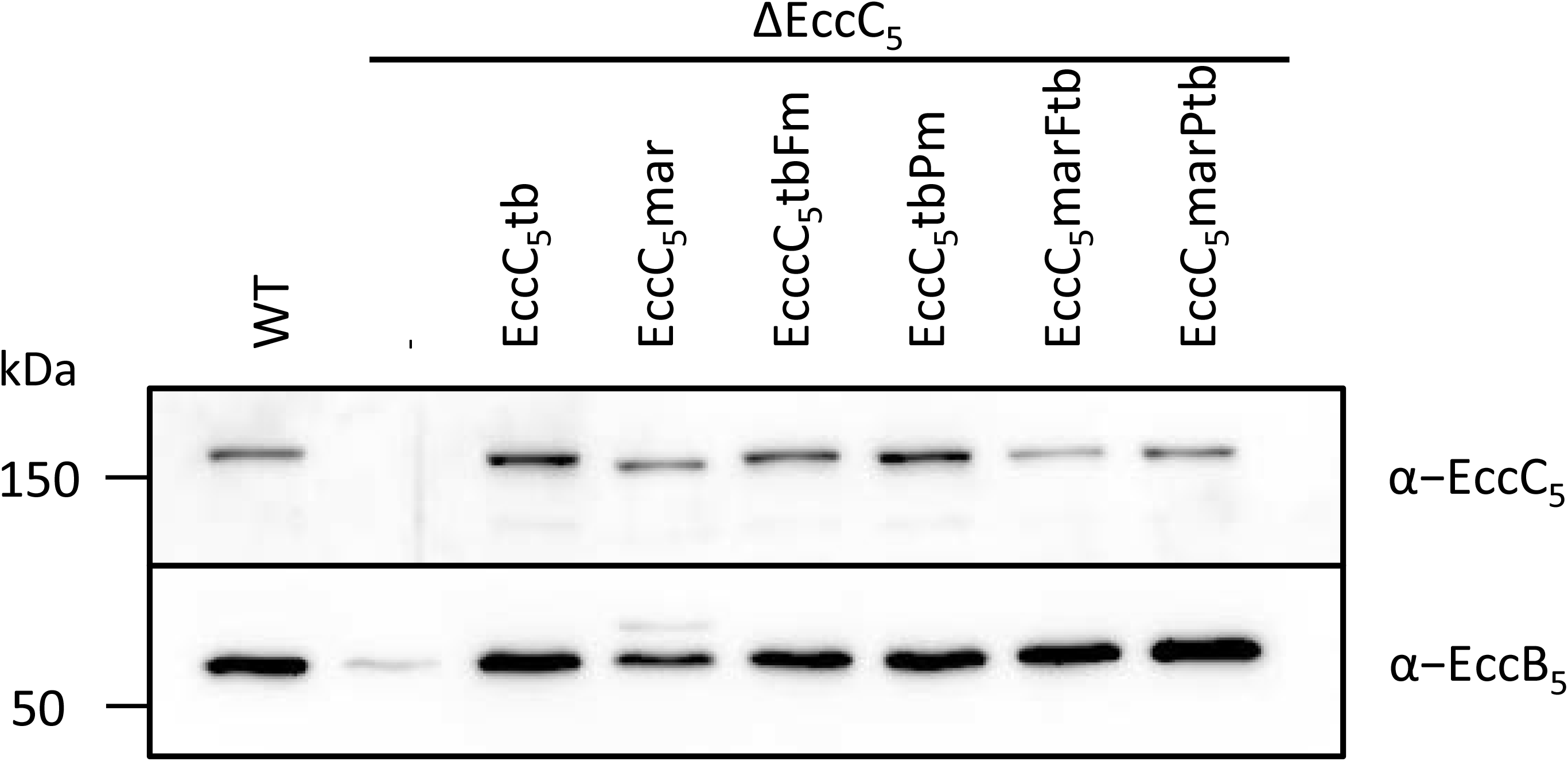
Expression of ESX-5 membrane components in *M. tuberculosis*. Immunoblot analysis of whole cells of *M. tuberculosis* WT, *ΔeccC_5_*and *ΔeccC_5_* complemented with constructs depicted in Fig. 2B and Fig S4. Proteins were visualized using antibodies against EccB_5_ and EccC_5_ (membrane components of ESX-5).

**Supplementary figure 6.**
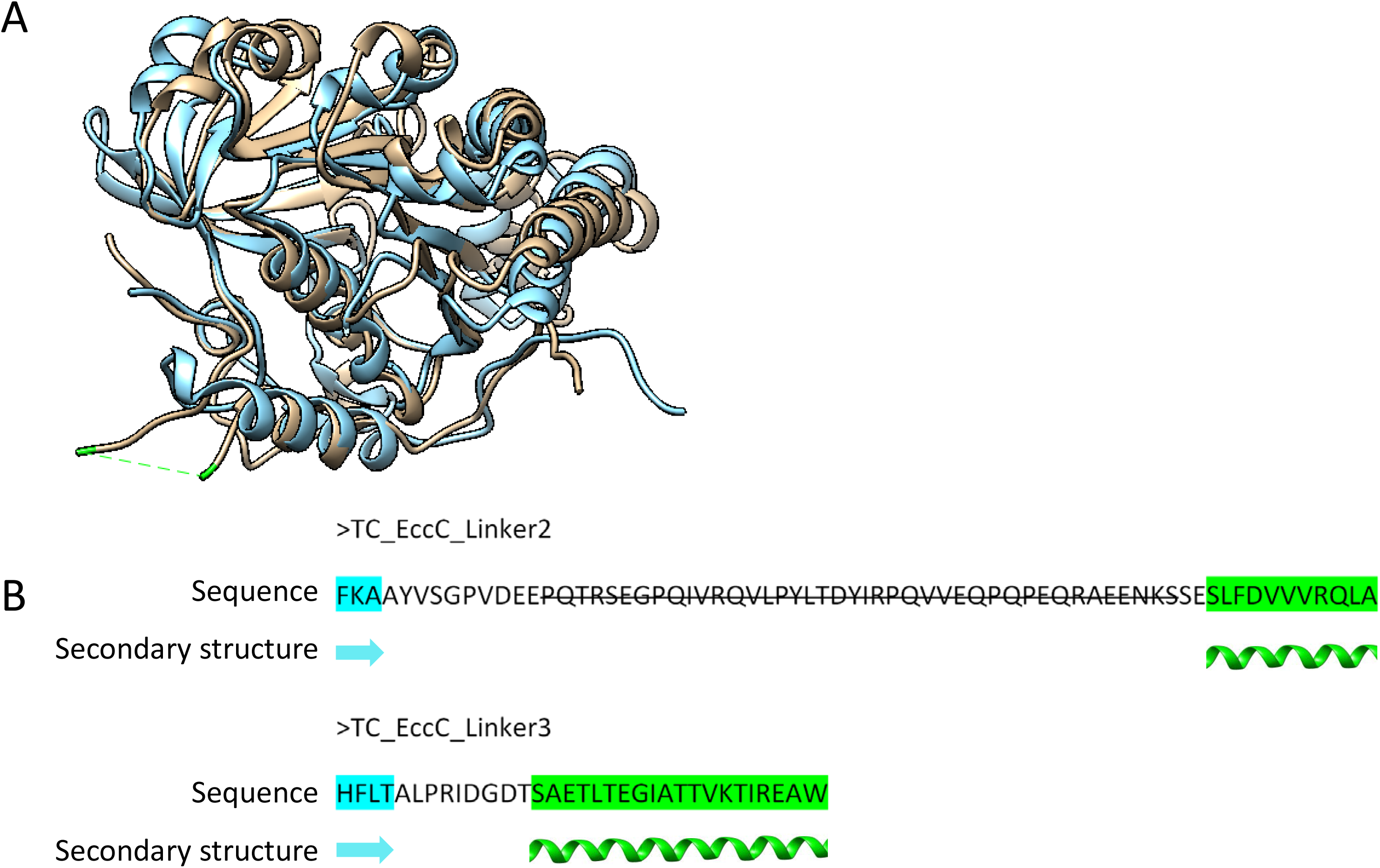
NBD1 and NBD2 superimposed. (A) NBD1 (light brown) and NBD2 (light blue) from *T. curvata* superimposed, emphasizing the missing structural feature in the linker 2 region, in green. (B) linker 2 and linker 3 loop sequence with adjacent upstream β-sheet (blue) and downstream α-helix (green). Striked through is the missing structural feature.

---

**Table S1.**
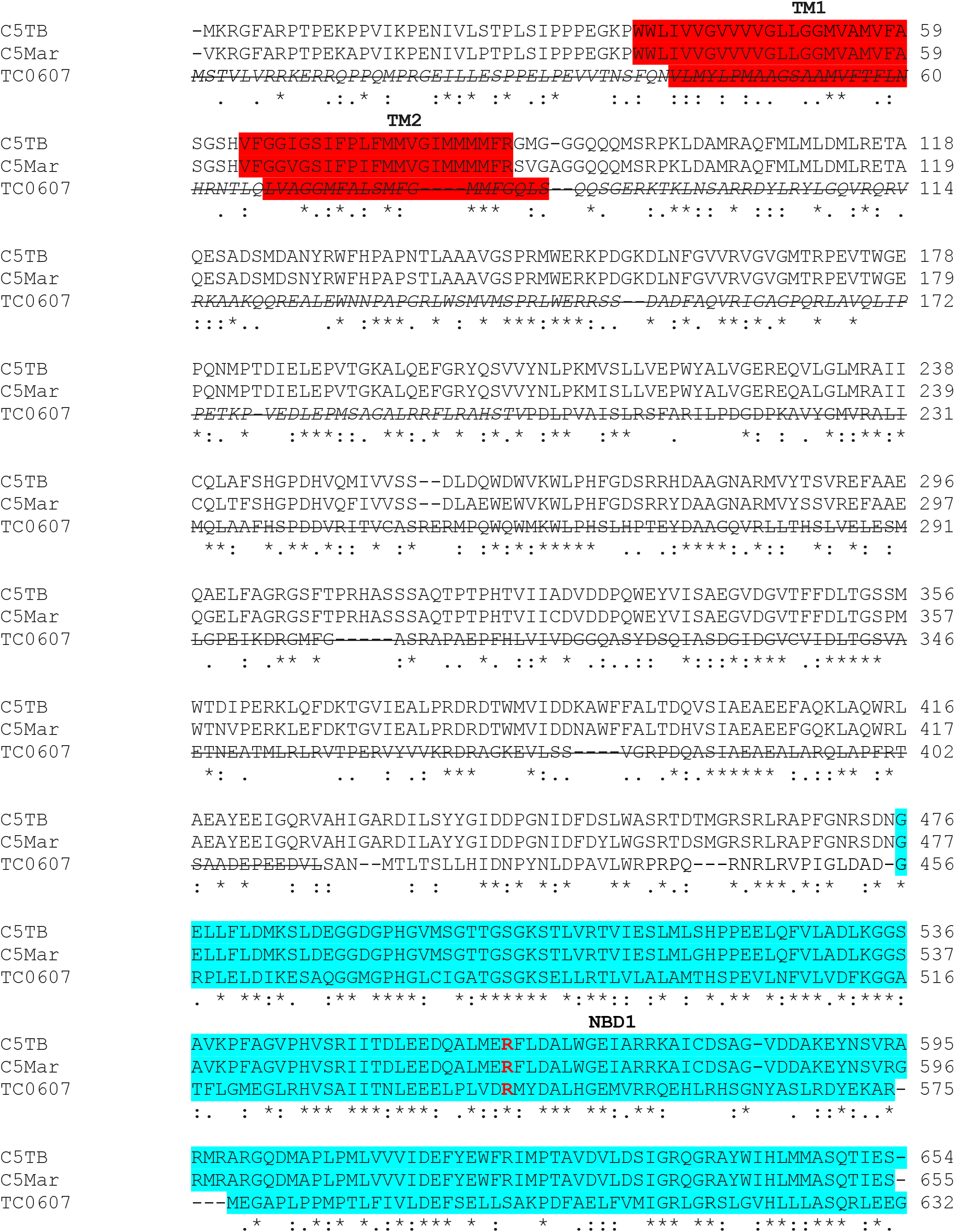
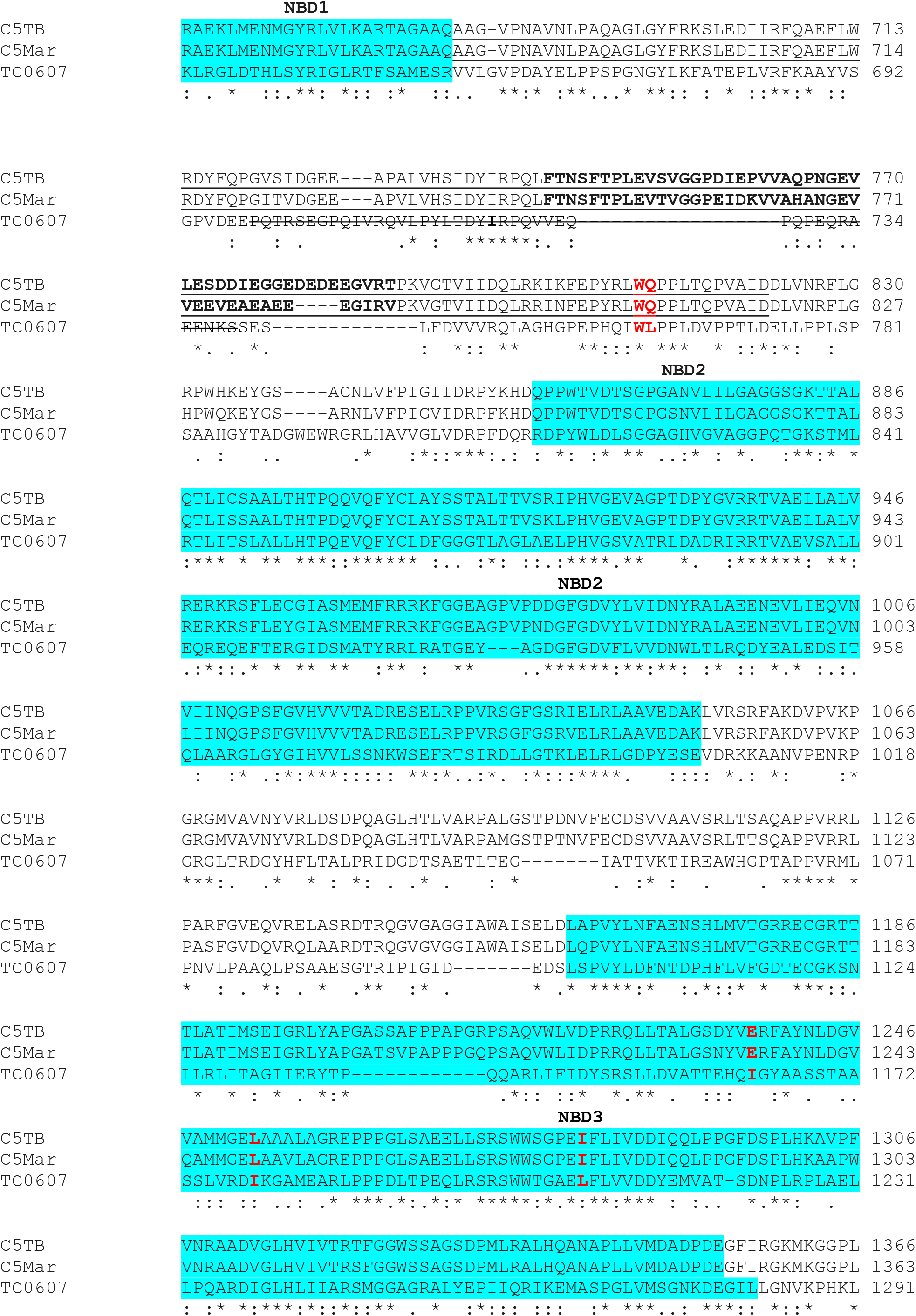
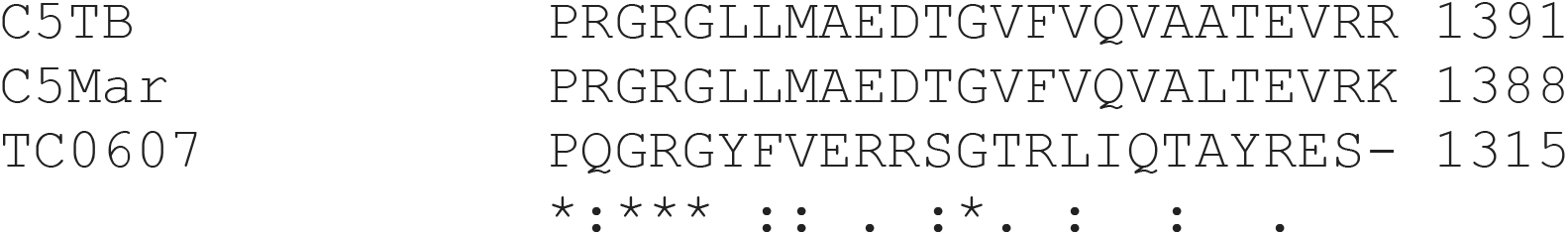
Alignment M. tuberculosis EccC_5_, M. marinum EccC_5_,T. curvata EccC.

**Table S2.**
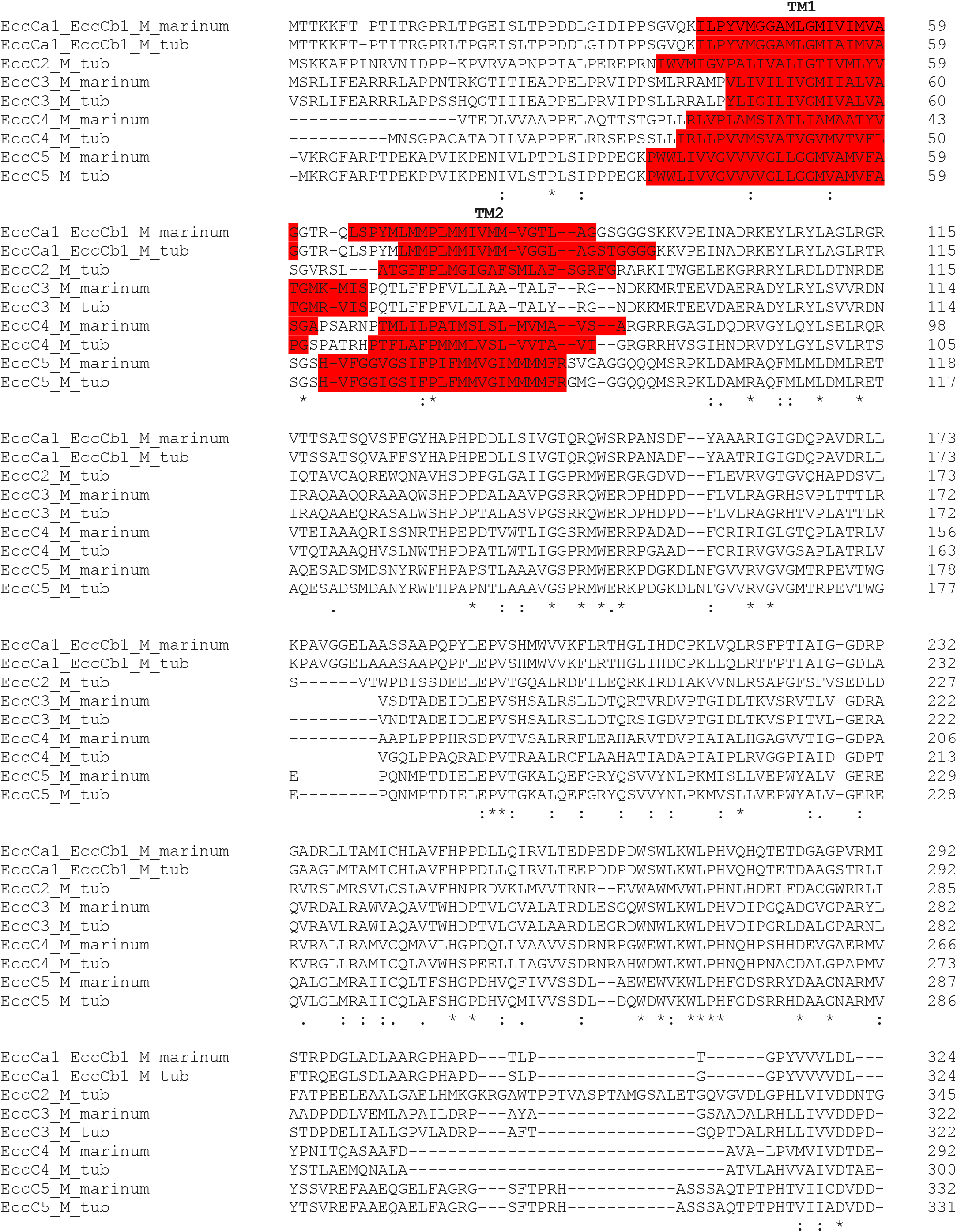
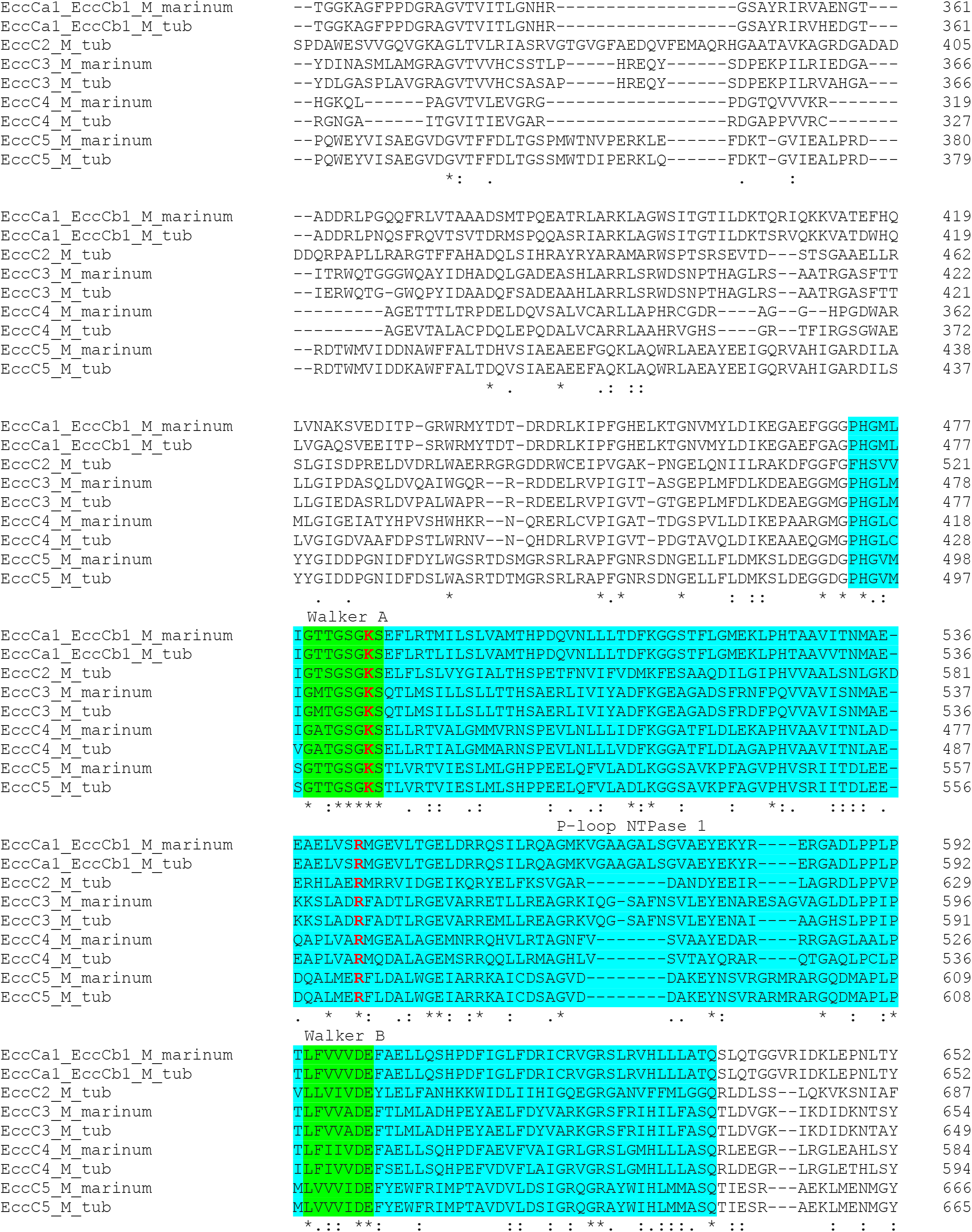
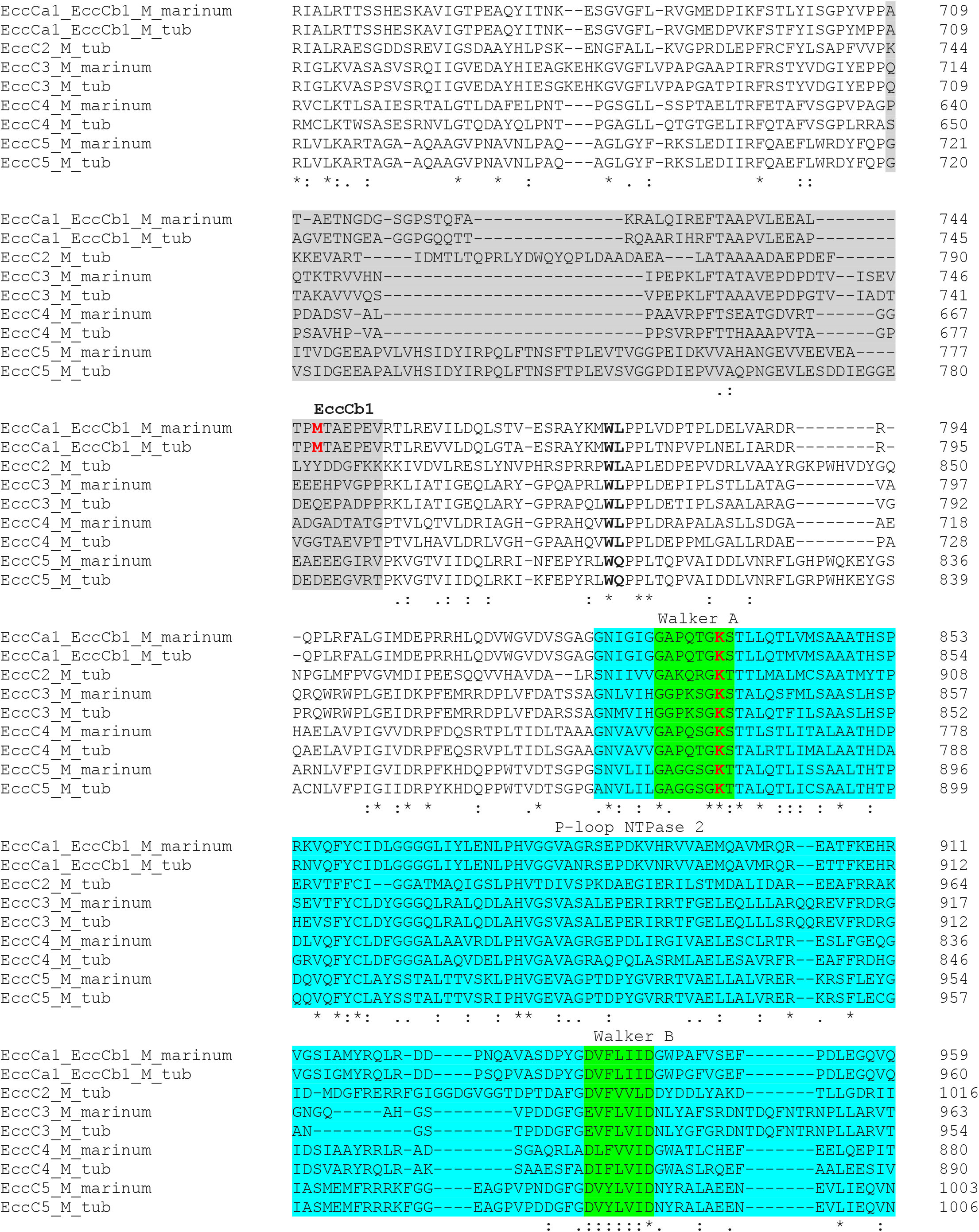
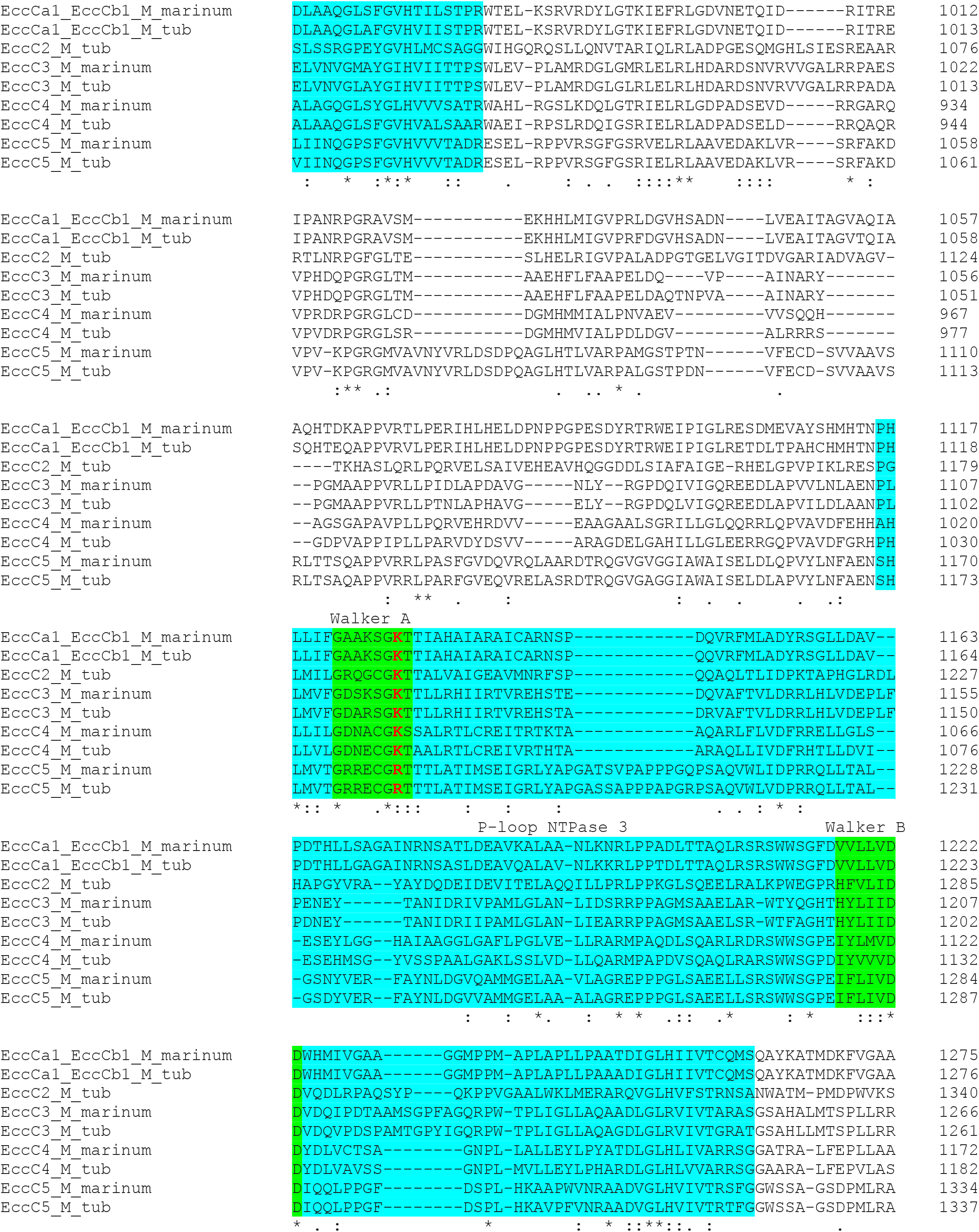
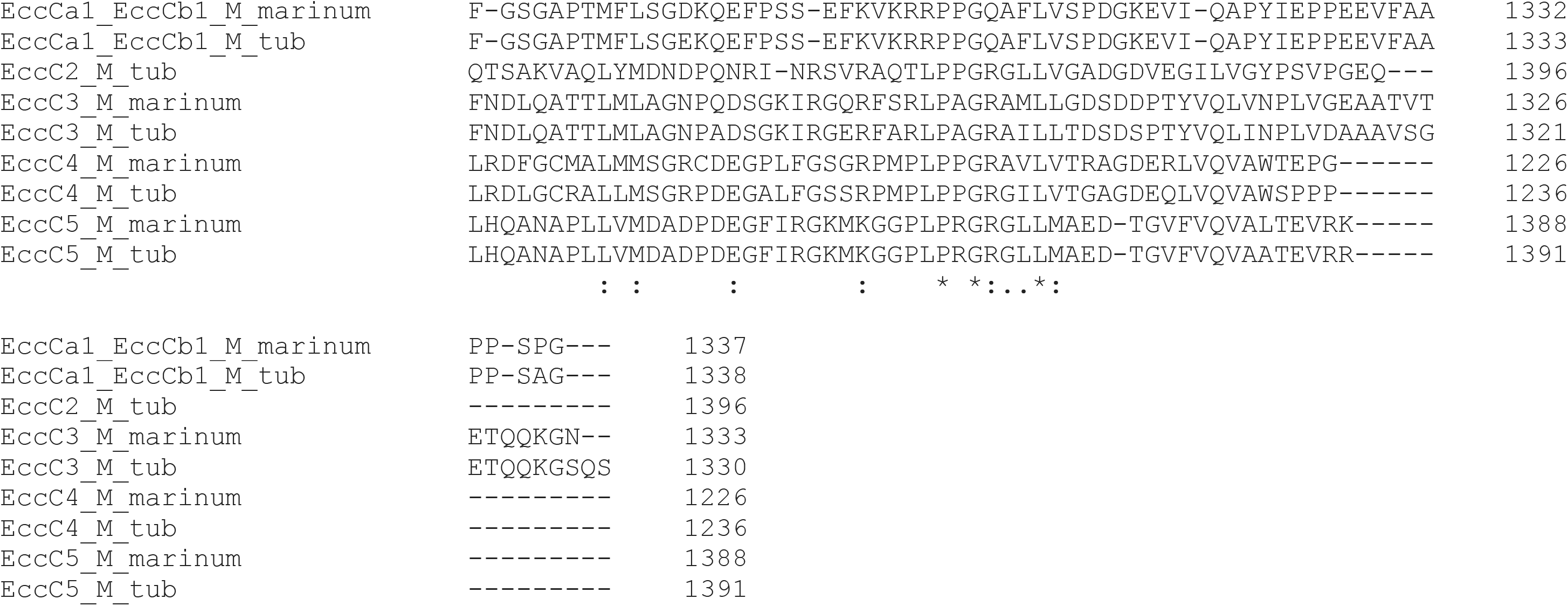
Alignment of all EccC homologs from M. marinum and M. tuberculosis.

**Table S3.** List of plasmids used in this study.

| Nr. | Plasmid | Features | Source |
| --- | --- | --- | --- |
| 1. | pMV-espG<sub>5</sub>-mtub | *hsp60* promotor, L5 *attP* & *aph*, *oriE*, *hyg*<sup>R</sup> | This study |
| 2. | pMV-espG<sub>5</sub>-mmar | *hsp60* promotor, L5 *attP* & *aph*, *oriE*, *hyg*<sup>R</sup> | Phan *et al.*, 2017 |
| 3. | pMV-eccC<sub>5</sub>-mtub | *hsp60* promotor, L5 *attP* & *aph*, *oriE*, *kan*<sup>R</sup> | This study |
| 4. | pMV-eccC<sub>5</sub>-mmar | *hsp60* promotor, L5 *attP* & *aph*, *oriE*, *kan*<sup>R</sup> | Ates *et al.*, 2015 |
| 5. | pMV-eccC<sub>5</sub>mtubF<sub>mmar</sub> | *hsp60* promotor, L5 *attP* & *aph*, *oriE*, *kan*<sup>R</sup> | This study |
| 6. | pMV-eccC<sub>5</sub>mtubP<sub>mmar</sub> | *hsp60* promotor, L5 *attP* & *aph*, *oriE*, *kan*<sup>R</sup> | This study |
| 7. | pMV-eccC<sub>5</sub>mmarF<sub>mtub</sub> | *hsp60* promotor, L5 *attP* & *aph*, *oriE*, *kan*<sup>R</sup> | This study |
| 8. | pMV-eccC<sub>5</sub>mmarP<sub>mtub</sub> | *hsp60* promotor, L5 *attP* & *aph*, *oriE*, *kan*<sup>R</sup> | This study |

**Table S4.**
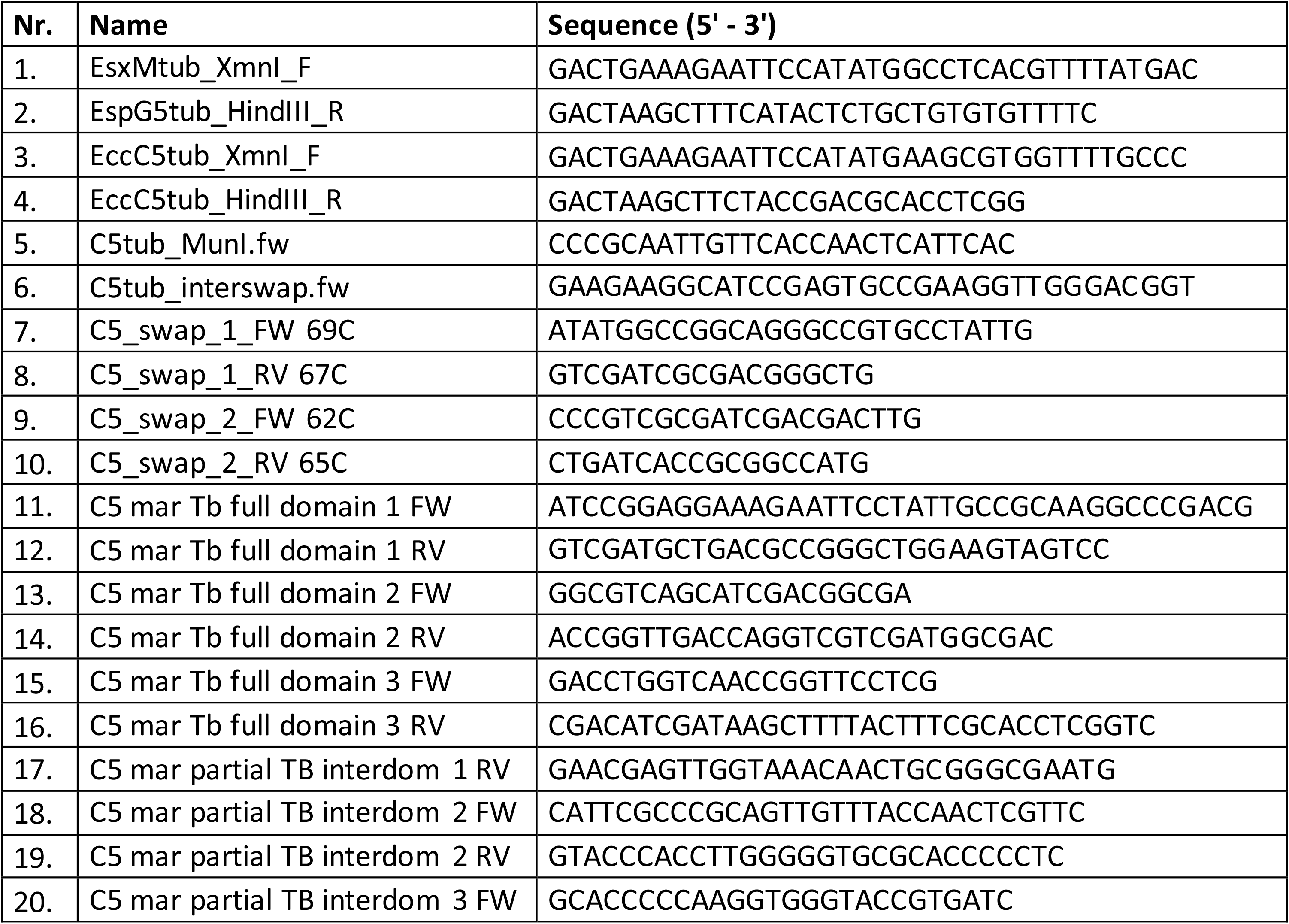
List of primers used in this study.

